# Replicating enzymatic activity by positioning active sites with synthetic protein scaffolds

**DOI:** 10.1101/2024.01.31.577620

**Authors:** Yujing Ding, Shanshan Zhang, Henry Hess, Xian Kong, Yifei Zhang

**Affiliations:** State Key Laboratory of Chemical Resources Engineering, Beijing University of Chemical Technology, Beijing 100029, China; Beijing Advanced Innovation Center for Soft Matter Science and Engineering, Beijing University of Chemical Technology, Beijing 100029, China; Department of Biomedical Engineering, Columbia University, 351L Engineering Terrace, 1210 Amsterdam Avenue, New York, NY 10027, United States; South China Advanced Institute for Soft Matter Science and Technology, Guangdong Provincial Key Laboratory of Functional and Intelligent Hybrid Materials and Devices, School of Emergent Soft Matter, South China University of Technology, Guangzhou 510640, P. R. China

## Abstract

Evolutionary constraints significantly limit the diversity of naturally occurring enzymes, thereby reducing the sequence repertoire available for enzyme discovery and engineering. Recent breakthroughs in protein structure prediction and *de novo* design, powered by artificial intelligence, now enable us to create enzymes with desired functions without relying on traditional genome mining. Here, we demonstrate a computational strategy for creating new-to-nature PET hydrolases by leveraging the known catalytic mechanisms and implementing multiple deep learning algorithms and molecular computations. This strategy includes the extraction of functional motifs from a template enzyme (here we use leaf-branch compost cutinase, LCC), regeneration of new protein scaffolds, computational screening, experimental validation, and sequence refinement. We successfully replicate PET hydrolytic activity with designer enzymes that are at least 30% shorter in sequence length than LCC. Among them, *Rs*PETase 1 stands out due to its robust expressibility. It exhibits comparable activity to *Is*PETase and considerable thermostability with a melting temperature of 56 °C, despite sharing only 34% sequence similarity with LCC. This work suggests that enzyme diversity can be expanded by recapitulating functional motifs with computationally built protein scaffolds, thus generating opportunities to acquire highly active and robust enzymes that do not exist in nature.

## Main

Enzymes are essential workhorse molecules of both living organisms and biochemical industries due to their powerful and sophisticated catalytic functions.^1-3^ The activity of an enzyme is often due to a small number of amino acid residues that are arranged in a defined three-dimensional configuration as a result of protein folding.^4^ This active site is surrounded by a large number of amino acids which serve as a protein scaffold holding the catalytic motifs in particular geometries. Similar catalytic functions are often found in enzymes that evolved independently with similar functional motifs from distinct ancestral proteins (examples of convergent evolution^5^), demonstrating that the protein scaffold can be varied. On the basis of this insight, past efforts of *de novo* design of enzymes have focused on the transplantation of functional motifs onto a foreign protein scaffold. Classic cases are the computational design of retro-aldolases^6^ and Kemp eliminase.^7^ A very recent advancement is the engineering of Fragaceatoxin C nanopores for hydrolyzing nano-sized polyethylene terephthalate (PET).^8^ However, this strategy relies on the availability of existing protein scaffolds with a certain level of structural similarity to the desired functional motifs.^9^ Furthermore, the introduction of new residues onto a native protein backbone may lead to unexpected geometrical variations in the active site.^10^ The limited availability of suitable scaffolds and the complexity of protein folding constrain our ability to design *de novo* enzymes.^11^

The rapid developments of artificial intelligence in protein science open up new opportunities to address these limitations. The deep learning program AlphaFold2 can predict the protein structure with atomic-level accuracy based solely on the primary amino acid sequence.^12, 13^ Two deep-learning methods, constrained hallucination and inpainting, can generate artificial protein backbones to scaffold pre-specified functional sites.^14^ Alternatively, a family-wide hallucination approach can generate large number of new idealized protein scaffolds for accommodating target substrates.^15^ A protein language model called ProGen was trained to produce protein sequences with a predictable function across diverse protein families.^16^ These advancements substantially improve the success rate for the design of new-to-nature enzymes.

The diversity of naturally occurring enzymes is constrained during evolution by the ancestral protein sequences and the need to maintain the homeostasis of a living system. *De novo* enzyme design increases the diversity of enzymes with desired activities without relying on conventional enzyme mining and discovery, which is usually labor-intensive and time-consuming.^17, 18^ For example, to address the global plastic pollution problem, researchers have made significant attempts to identify potential PET hydrolytic enzymes from the environment for two decades.^19, 20^ Even though these efforts have resulted in dozens of different enzymes with known activity against PET,^21^ so far the most promising ones for industrial PET recycling are *Ideonella sakaiensis* PETase (*Is*PETase, 290 amino acids, M_w_=30.25 kDa)^20, 22^ and leaf-branch compost cutinase (LCC, 258 amino acids, M_w_ = 27.78 kDa)^23^. These enzymes share a conserved catalytic triad of serine (Ser)-histidine (His)-aspartate (Asp) residues borne by distinctly different protein backbones. This inspired us to design new-to-nature PET hydrolases by re-scaffolding the catalytic triad with synthetic protein scaffolds.

Our approach to achieve PET hydrolytic activity with designer enzymes relied on a combination of computational and experimental techniques. Building on the demonstration that the inpainting method has the ability to generate new proteins with binding activity inherent in functional motifs,^14^ we employed an inpainting approach to generate new sequences encoding protein scaffolds to support the catalytic triad of LCC and some adjacent residues. Taking advantage of the protein structures predicted by ColabFold (an online tool integrating AlphaFold2 with the fast homology search of MMseqs2),^24^ we conducted *in silico* screening for the virtual designer enzymes that are putatively capable of hydrolyzing PET. We evaluated the expression and function of 10 designer enzymes obtained from the screening, and accordingly revisited and refined the sequences by iterative computation. Two designer enzymes were obtained that exhibit PET hydrolytic activity but have low expressibility. After sequence refinement using ProteinMPNN,^25^ we created an expressible novel PET hydrolase with PET hydrolytic activity comparable to the wild-type *Is*PETase.

The successful creation of a synthetic scaffold for a known catalytic mechanism demonstrates the effectiveness of deep learning methods in the construction of designer enzymes. By computationally exploring the protein universe without relying on conventional genome mining strategies, we are now able to create new-to-nature enzymes with specific activities, ushering in a new age of enzyme design.

### A general workflow for the computational design of PET hydrolases

Currently known PET hydrolases, including LCC and *Is*PETase, possess the same catalytic triad of Ser-His-Asp. The specific spatial arrangement of these key amino acids endows the ester hydrolytic function, at the same time the amino acids around the catalytic triad form a substrate-binding groove that helps to stabilize the reaction transition state via electrostatic interactions and hydrogen bonding.^26, 27^ Since LCC and its variations are more catalytically active and thermally stable than *Is*PETase, we attempt to generate new-to-nature PET hydrolases by re-scaffolding the functional motifs of LCC. Moreover, as LCC is the smallest PET hydrolase yet discovered, we sought to design an even smaller designer counterpart, which could potentially improve the cost-effectiveness of industrial processes by reducing the amount of enzymes used. The workflow we follow to generate designer enzymes is shown in a simplified version in Fig. 1. We first identify the active sites of LCC based on molecular docking, and then extract the catalytic motifs together with adjacent pieces of secondary structures. We use RF_joint_-guided inpainting to fill in additional sequences encoding new protein structures to scaffold the extracted structures.^14^ We then use ColabFold to predict the three-dimensional protein structures of the generated sequences. The virtual structures with high prediction certainty are selected for further computational screening. The screening considers the motif root mean square deviation (RMSD) between the backbone atoms of the retained LCC segments and those from the virtual enzymes, the binding energy with a substrate molecule, and the time-dependent Cα-RMSD by Molecular Dynamics (MD) simulation. We then express the screened virtual enzymes in *E. coli* cells and characterize the enzymatic PET hydrolytic activity. The problematic sequences can be revised by executing the RF_joint_ step again and the active but difficult-to-express sequences can be rescued by using ProteinMPNN. By iteratively performing the scaffold remodeling, computational screening, experimental validation and sequence refinement, we can obtain active and expressible new enzymes whose sequence is very different from the template. This workflow is in principle applicable to the de novo design of other enzymes with known catalytic motifs.

**Fig. 1.**
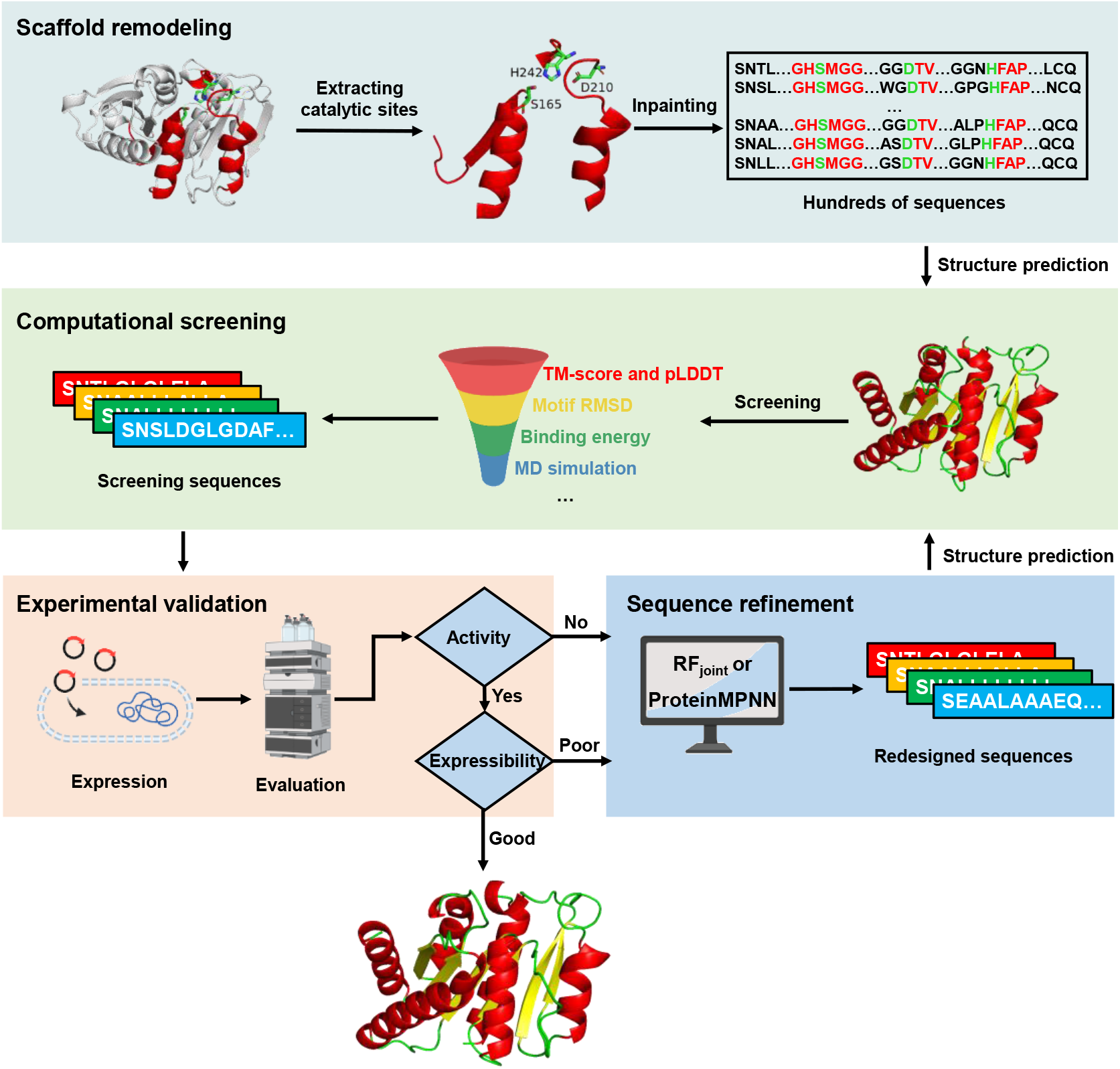
A workflow for the computational design of new-to-nature PET hydrolases by remodeling the protein scaffold. The workflow includes four steps: scaffold remodeling, in which the functional motifs are extracted from a template enzyme and then the missing sequences are completed by inpainting; computational screening, in which the newly generated sequences are screened computationally based on a set of physiochemical criteria; experimental validation, in which the filtered sequences are examined in terms of expression and expected activity; sequence refinement, in which flaws in designs are revised. Iterative implementation of the sequence refinement, computational screening, and experimental validation steps effectively improves the quality of designs.

### In silico generation and screening of virtual PET hydrolases

To guarantee the success rate of the computational design and minimize the search space, we retained three disconnected LCC sequence segments so that each segment encodes a structural motif containing a key residue of the indispensable catalytic triad, S165/D210/H242. The retained sequences consist of 44 amino acid residues in total, accounting for 17% of the full-length LCC. We then generate complementary protein scaffolds to recapitulate the retained motifs in place by using a RF_joint_-guided inpainting approach. We intentionally restricted the length of the *in silico* generated sequences to be shorter than LCC (see Supplementary Table 1). The inpainting step generates hundreds of distinct protein sequences ranging in length from 140 to 185 aa within only a few minutes. The three-dimensional protein structures of these sequences were predicted by ColabFold, which offers fast structure prediction based on AlphaFold2 and an orthogonal validation of the putative folding of sequences generated by the RF_joint_. The quality of the prediction can be evaluated by the per-residue confidence score predicted LDDT (pLDDT) (estimating how well the prediction would agree with an experimental structure) and template modeling scores (measuring the similarity of the predicted structure to a known template). The predictions with a pLDDT > 70 and a pTM score > 0.7 were kept for further screening. In order to further narrow down the candidate pool of the virtual enzymes that are potentially active, we screened based on the accuracy of motif recapitulation, the interactions between the model substrate and the virtual enzymes, and the structural stability during MD simulations. The accuracy of motif recapitulation was assessed by the motif RMSD between the original motifs from LCC and those from the virtual enzymes. The virtual enzymes with motif RMSD larger than 2.5 Å were discarded.^28^ Fig. 2a shows a 3D scatter plot of 75 designs, of which 44 meet these static structural quality criteria (blue dots). The right upper panel depicts a typical structural superposition of the retained segments in a predicted protein structure against the template, where the motif RMSD is only 2.0 Å, indicating a successful motif recapitulation. The filtered virtual enzymes were further examined via molecular docking with the model substrate 2-HE(MHET)_3_. A typical molecular docking of the model substrate 2-HE(MHET)_3_ with a virtual enzyme is shown in Fig. 2b. As the binding energy between LCC and 2-HE(MHET)_3_ was calculated to be -2.78 kcal/mol, we chose a range of -4 to 0 kcal/mol as a screening criterion. Furthermore, the reaction mechanism necessitates two criteria: the distance between the carbonyl carbon atom of 2-HE(MHET)_3_ and the oxygen atom of serine (Ser) (< 4 Å), and the potential to form an oxyanion hole by the backbone NH groups.^29^ The structural stability of the virtual enzymes was assessed by calculating the Cα-RMSD evolution through MD simulations. The virtual enzymes whose RMSD values exceed 5 Å within an 18-ns simulation were discarded (Fig. 2c). Ten virtual enzymes (designated as P1 through P10) out of 100 candidates passed the computational screening steps described above. The molecular docking of these enzymes with 2-HE(MHET)_3_ was shown in Supplementary Fig. 1. All these enzymes are shorter in length than LCC, with full lengths between 140 and 186 and molecular weights between 14.28 kDa and 19.31 kDa. Detailed descriptions including their sequences, lengths, and molecular weights are summarized in Supplementary Table 1, and the predicted protein structures are shown in Supplementary Fig. 2.

### Expression and experimental characterization of designer PET hydrolases

The ten virtual enzymes were expressed in *E. coli* BL21 (DE3) strains. The enzymes were fused with His-tags at N-terminals, allowing ease purification by Ni-NTA chromatography (Fig. 2d). The crude proteins eluted from the column were analyzed by sodium dodecyl sulfate-polyacrylamide gel electrophoresis (SDS-PAGE) (Supplementary Fig. 3). We did not see the formation of inclusion bodies in the cell lysates. All eluting protein fractions contain only a small fraction of the target enzyme in the presence of a substantial amount of protein impurities, showing that the target proteins were poorly expressed (less than 100 ng/mL in the cell lysates). The low expression levels indicate a low quality of the *in silico* designed sequences. Nevertheless, we tested the activity of these enzymes in the protein mixture. P1, P3, P4, P5, P7, P8 and P9 are capable of hydrolyzing bis(2-Hydroxyethyl) terephthalate (BHET) into mono(2-hydroxyethyl) terephthalate (MHET) and terephthalate (TPA), suggesting that these designer enzymes are esterases (Fig. 2e). However, none of these enzymes exhibit any PET hydrolyzing activity, indicating that the actual folding of these virtual enzymes may differ from what is predicted.

**Fig. 2.**
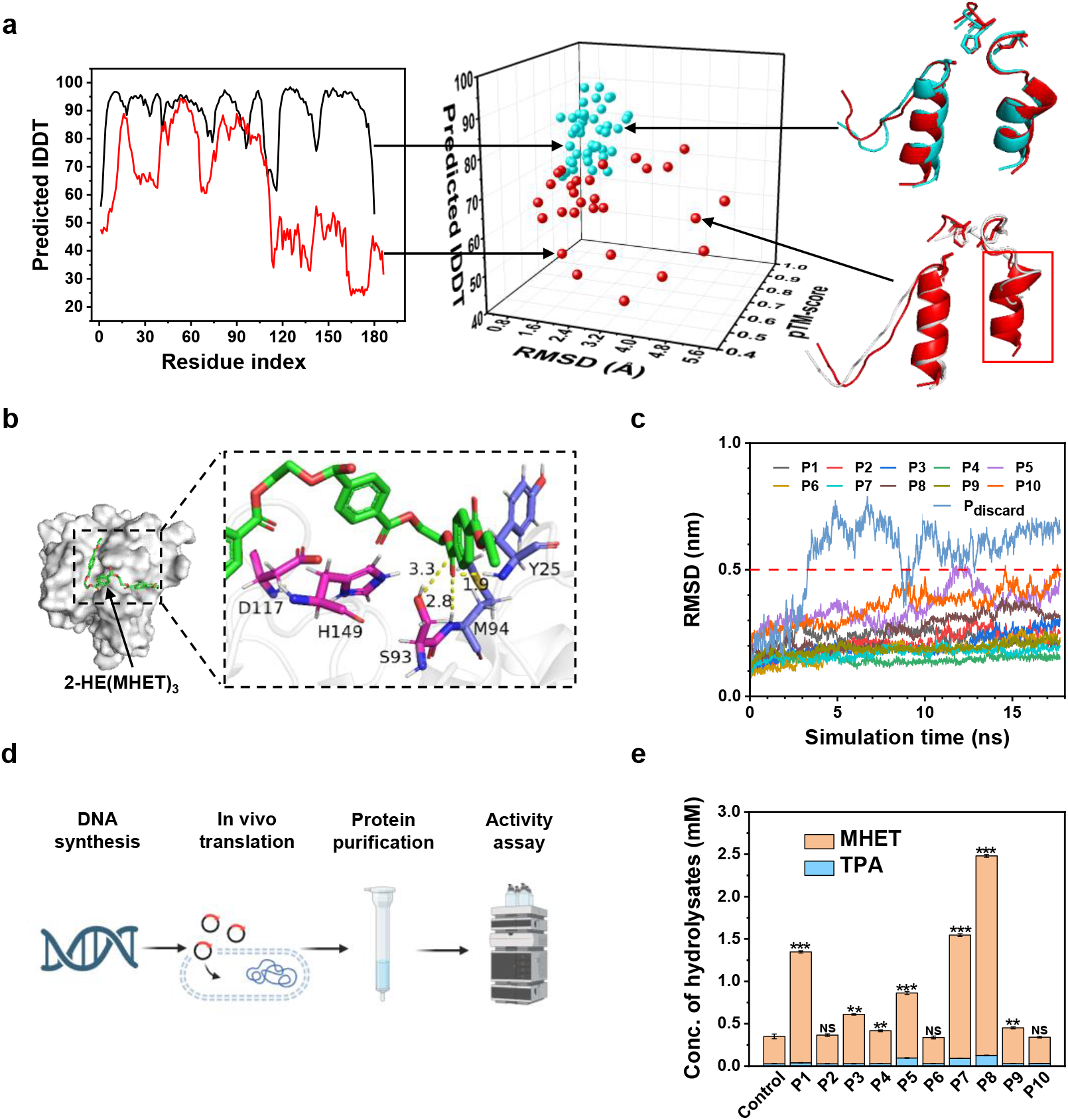
Typical results of the computational screening. **a**, A 3D scatter plot showing the AlphaFold pLDDT, pTM-score and the motif RMSD of the computationally generated virtual enzymes. Examples of virtual enzymes with qualified pLDTT (black curve) and unqualified pLDTT (red curve) are shown on the left. Examples of virtual enzymes with the motif RMSD of 2.00 Å (blue) and 4.82 Å (white) compared to the retained motifs from LCC (red) are shown on the right. **b**, Molecular docking of 2-HE(MHET)_3_ (green stick model) in a virtual enzyme. 2-HE(MHET)_3_ interacts with the catalytic triad (magenta stick model) and the oxyanion hole potentially formed by the NH groups of Y25 and M94. **c**, Time-course RMSD of Cα of representative virtual enzymes assessed by MD simulations. **d**, Procedures for the expression, purification and activity assay of the designer enzymes. **e**, The hydrolysis of BHET by the eluted protein fractions from a Ni-NTA affinity column. The reaction system contained 6.0 μg/mL proteins and 1.0 mg/mL BHET and was incubated at 25 °C for 24 h. BHET hydrolyzing activity of P1-10 was compared with the control, the self-hydrolysis of BHET in the same condition, using a t-Test, P <0.001 is denoted by^***^, P <0.01 is denoted by^**^, P <0.05 is denoted by^*^, and P >0.05 is denoted by NS (no significant difference). Error bars represent the standard deviations of three measurements.

### Rescuing the designs by iteratively running RFjoint

We assume that the poor expressibility and the absence of PET hydrolytic activity are due to the inherent flaws in the sequences designed in the first round. A review of the sequences revealed that the RF_joint_ algorithm tends to produce long single amino acid repeats (SAAR) such as LLLLLLL and GGGGGGGG, and consecutive hydrophobic amino acids (CHAA) such as LVVLV, which are uncommon in native protein sequences (Supplementary Table 2). The predicted protein structures are shown in Supplementary Fig. 4. The lack of amino acid diversity in the generated sequences has been discussed by Zheng *et al*. as an inherent limitation of structure-based protein sequence design models.^30^ In the predicted 3D structures of virtual proteins, we found large regions of hydrophobic patches on the protein surface, which are most likely the source of protein misfolding and aggregation.^31^ These regions may be identified as aberrant by the cellular proteolytic machineries, such as the Lon protease, leading to a rapid degradation of the as-synthesized peptides.^32^ Therefore, we iteratively employ RF_joint_ to generate new sequences to replace the problematic ones encoding long SAAR and CHAA patterns and large hydrophobic patches (Fig. 3a). Typically, several runs of iterations were enough to replace the problematic areas to the maximum degree that RF_joint_ can repair. The redesigned sequences were filtered by the aforementioned computational screening steps. We eventually obtained three new designer enzymes, namely P4-a (derived from P4), P5-a (derived from P5) and P7-a (derived from P7). The surface hydrophobic area of the designer enzymes (Supplementary Table 3) significantly decreases after sequence refinement, making the designs more akin to natural proteins. The regional pLDDT values of the redesigned sequences as well as the overall pLDDT and pTM-scores of the full-length sequences are significantly improved (Supplementary Fig. 5 and Supplementary Table 1), indicating a higher structural quality predicted by ColabFold. The theoretical molecular weights of these enzymes were 19.43 kDa, 18.47 kDa and 18.34 kDa, respectively, which are 30–34% smaller than the molecular weight of LCC. The three designer enzymes share only 47%, 46%, and 47% sequence similarities compared to LCC (right panel of Fig. 3a and Supplementary Fig. 6). Nevertheless, the molecular docking of these enzymes with 2-HE(MHET)_3_ show that the substrate molecule is accommodated in the active site with appropriate interactions for catalysis (Supplementary Fig. 7).

**Fig. 3.**
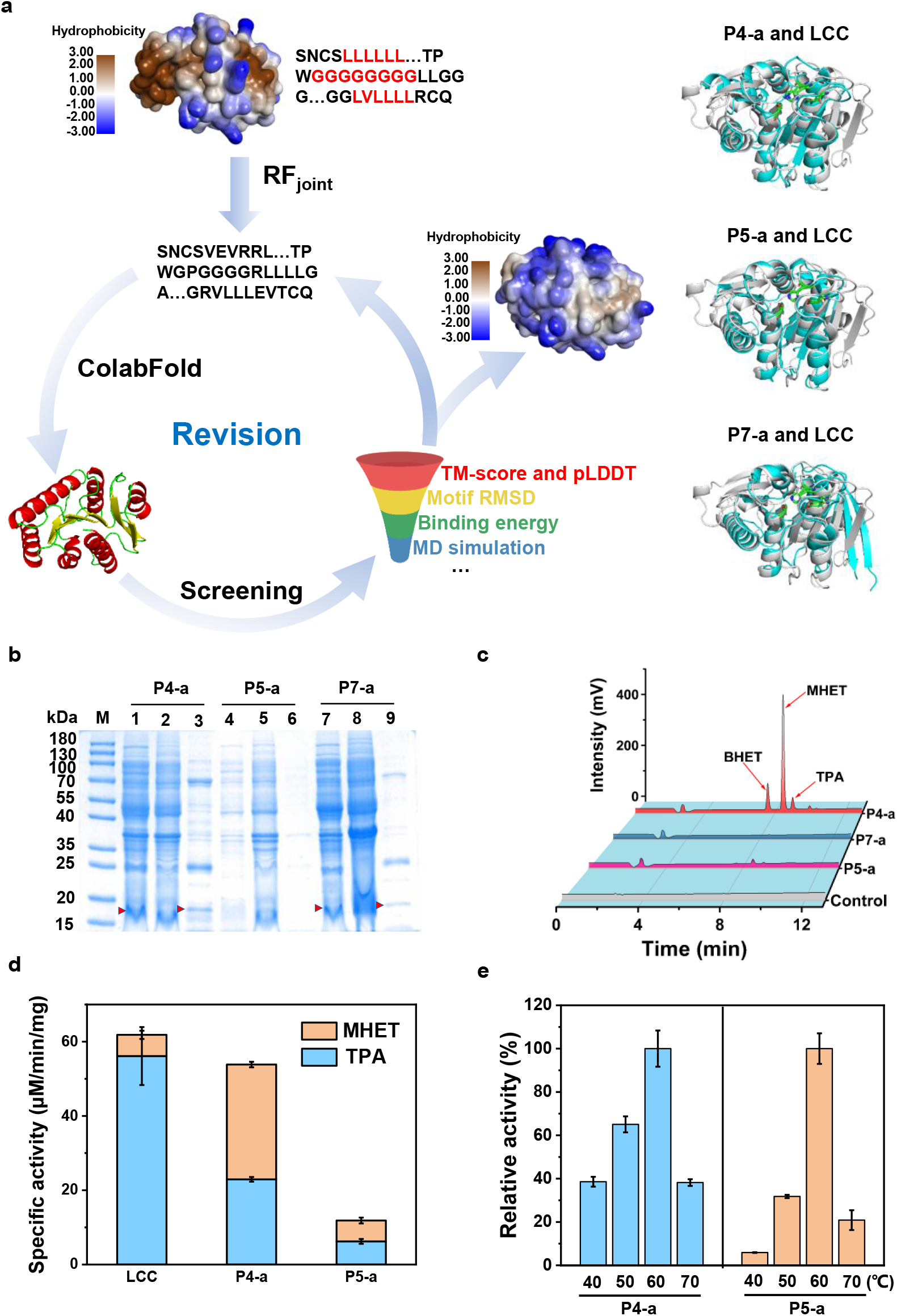
Computational refinement of the designed sequences. **a**, The exposed hydrophobic regions on the protein surface (brown) and the amino acid sequences of SAAR and CHAA patterns (red) were redesigned by RF_joint_, followed by 3D structure prediction and computational screening. The right panel shows the structural comparisons between P4-a, P5-a, P7-a (white) and the template enzyme LCC (blue), respectively. **b**, SDS-PAGE analysis of P4-a, P5-a and P7-a. Lanes 1, 4, 7: P4-a, P5-a and P7-a in the soluble fraction of the cell lysate; Lanes 2, 5, 8: P4-a, P5-a and P7-a in the precipitates of the cell lysate; Lanes 3, 6, 9: P4-a, P5-a and P7-a in the eluted fraction after Ni-NTA affinity chromatography. **c**, Hydrolysis of PET by P4-a, P5-a and P7-a at 50 °C for 24 h. The concentrations of hydrolysates BHET, MHET and TPA were determined by HPLC. **d**, Specific activities of LCC, P4-a and P5-a against PET. **e**, The activity-temperature dependence of P4-a and P5-a. All assays were carried out with 5 mg/ml of low-crystallinity PET powder in sodium phosphate buffer (50 mM, pH 8.0). Error bars represent the standard deviations of three measurements.

We then expressed P4-a, P5-a, and P7-a in Lemo21 (DE3) competent *E. coli* strains, a BL21(DE3) derivative harboring the pLemo plasmid, which is suitable for the expression of challenging proteins.^33^ After the expression and purification, the eluting proteins were subjected to SDS-PAGE analysis (Fig. 3b). Unfortunately, the expression of these enzymes did not improve substantially, and the band for P5-a was almost invisible. We determined the PET hydrolytic activity of these new designer enzymes (in the eluted fraction from Ni-NTA column) using low-crystalline PET powder as the substrate. Both P4-a and P5-a exhibit PET hydrolytic activity, as evidenced by the release of PET hydrolysates MHET and TPA (Fig. 3c).

### Enhancing the protein expression using SUMO fusions

The expression problem has occurred in other PET hydrolases, such as the one mined from the human saliva metagenom.^19^ Fusion to SUMO (small ubiquitin-like modifier) tags is considered effective in boosting the expression level of recombinant proteins by improving folding, solubility, and stability.^34^ We therefore expressed SUMO-tagged P4-a and P5-a in *E. coli* BL21 (DE3) PLysS cells. However, SUMO-fusion did not substantially improve the expression of these two enzymes, and their concentrations in cell lysates were still insufficient for purification (Supplementary Fig. 8a). Through quantitative western blotting, we identified the presence of P4-a and P5-a and quantified their concentrations to be 1.7 and 2.9 μM after Ni-NTA column purification, respectively (Supplementary Fig. 8b, c). The SUMO tags on P4-a and P5-a were then removed by adding SUMO-specific proteases ULP1. We determined the specific activity of P4-a and P5-a toward the low-crystallinity PET powders and found that P4-a exhibits higher PET hydrolytic activity than P5-a. P4-a has comparable PET degradation activity to the original template LCC but accumulates more MHET, suggesting that it has lower activity toward MHET (Fig. 3d). The temperature optima of P4-a and P5-a (Fig. 3e) are both 60 °C, demonstrating that these two designer enzymes have excellent thermostability.

### Rescuing the designs using ProteinMPNN algorithm

The failure in expression of the above enzymes indicates the presence of elusive imperfections of the computationally generated sequences, which is a common issue of *de novo* designed proteins.^35^ Dauparas *et al*. has recently demonstrated that ProteinMPNN can rescue some failed designs created by Rosetta or AlphaFold, and the redesigned sequences showed increased solubility and thermostability.^25^ Encouraged by this, we employed ProteinMPNN to optimize the sequences of the potentially active designer enzymes P4-a and P5-a based on their respective backbones. We generated 10 new sequences for each backbone and predicted their 3D structures using ColabFold. After *in silico* screening with the above-mentioned computational approaches, we finally selected 4 sequences for expression in *E. coli* BL21 (DE3) strains, namely P4-a-1 and P4-a-2 (derived from P4-a), P5-a-1 and P5-a-2 (derived from P5-a). The backbone RMSD values of these proteins before and after optimization by ProteinMPNN are less than 1.0 Å (Fig. 4a), and the optimized structures exhibit increased structural quality and stability (Supplementary Figs 9 and 10). All redesigned enzymes exhibited hydrolytic activities toward BHET (Supplementary Fig. 11) and low-crystallinity PET powder (Fig. 4b). Only P4-a-2 was successfully expressed and isolated with 95% purity, and the expression level was estimated to be 2.5 mg/L (Fig. 4c, d). The other three proteins, however, were almost invisible in SDS-PAGE analysis (Supplementary Fig. 12a, b). Most impressively, the temperature optimum of P4-a-2 for PET hydrolysis is 50 °C (Fig. 4e). This is in line with its melting temperature (T_m_ = 56 °C) characterized by the differential scanning fluorimetry (DSF) assay (Supplementary Fig. 13), suggesting that P4-a-2 has good thermostability. The catalytic efficiency of P4-a-2 on PET was 0.5 L/g/s (Fig. 4f), comparable to that of wild-type *Is*PETase.^36^ Pairwise sequence alignment shows that P4-a-2 shares only 34% sequence similarity to LCC and 25% to *Is*PETase (Supplementary Fig. 14). This implies that the catalytic function can be maintained even if a significant portion of protein sequences is altered by our strategy. As this is the first designer PETase designed by computational *Re-scaffolding*, we refer to it as *Rs*PETase 1. Since the isolation of *Is*PETase in 2016,^20^ continuous efforts have been devoted to improve its activity. The most recent reported mutants, HotPETase^37^ and FAST-PETase^38^, have three orders of magnitude increased PET hydrolytic activity compared to the wild-type.^39, 40^ Based on these achievements, we believe that the PET hydrolytic activity of *Rs*PETase 1 can also be significantly improved after several rounds of protein engineering.

**Fig. 4.**
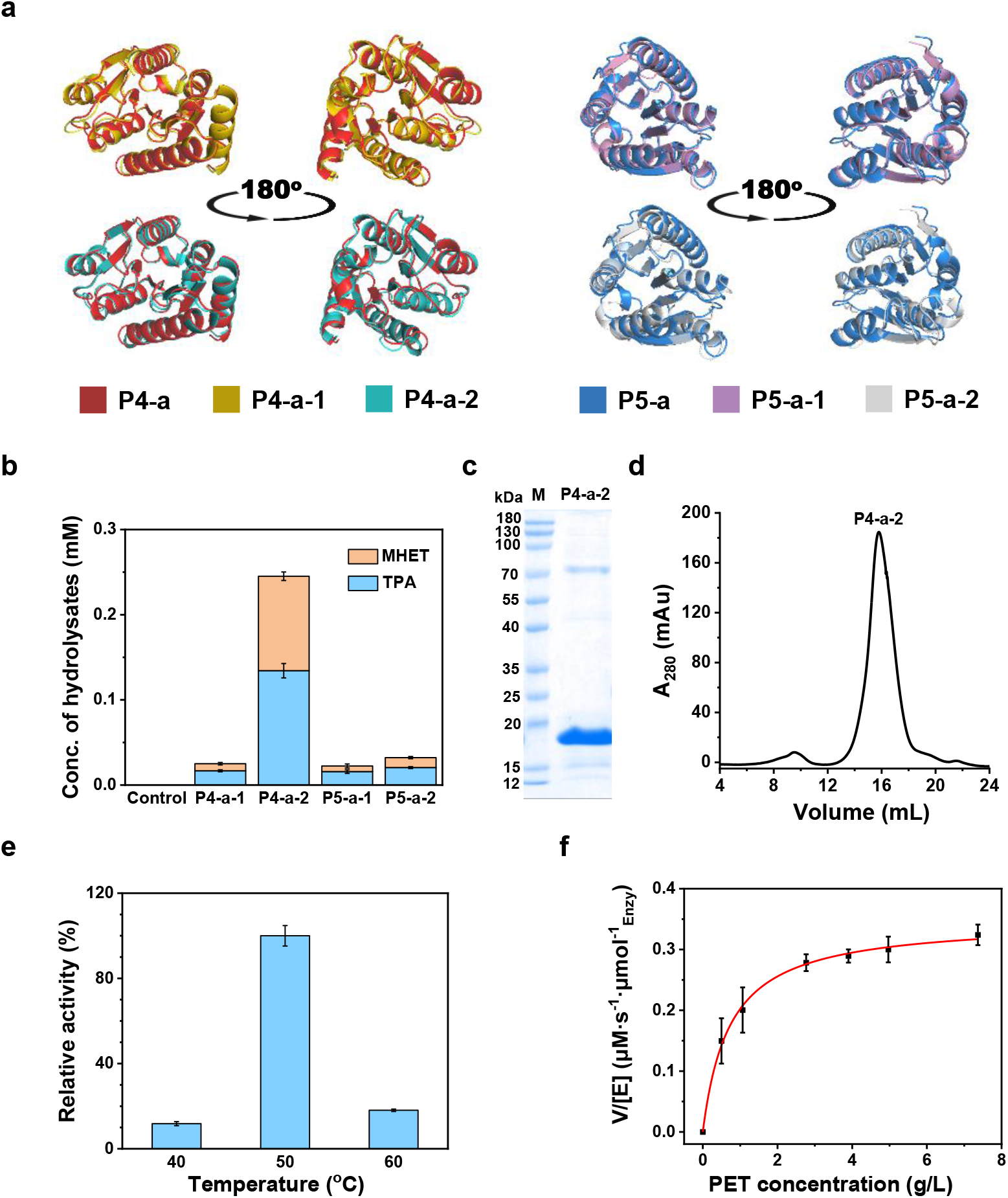
Expression and activity assays of ProteinMPNN-rescued P4-a and P5-a. **a**, Superimposition of the 3D structures of redesigned proteins on the corresponding original protein. P4-a-1 and P4-a-2 (also named *Rs*PETase 1) are redesigned from P4-a; P5-a-1 and P5-a-2 are redesigned from P5-a. Protein structures were predicted by ColabFold. The backbone RMSDs for every pair of backbones (from left to right) are 0.58 Å, 0.57 Å, 0.69 Å, and 0.82 Å, respectively. **b**, Concentrations of hydrolysates released by the redesigned enzymes. Reactions were carried out with 6.0 μg/mL proteins in the eluted fraction and 5.0 mg/mL PET powder at 50 °C for 24 h. **c**, Coomassie-stained SDS-PAGE analysis of *Rs*PETase 1. **d**, Size-exclusion chromatography of *Rs*PETase 1. **e**, Temperature dependence of *Rs*PETase 1. **f**, Michaelis-Menten plot of *Rs*PETase 1.

## Discussion

We have demonstrated a successful workflow for the computational design of new-to-nature PET hydrolases by scaffolding the known active sites using a combination of recently published deep learning algorithms. A set of physiochemical criteria was established for computational screening of putatively foldable and active designer enzymes. The filtered designs were validated by expression in *E. coli*. cells. Failed designs were rescued by iterative inpainting using RF_joint_ and sequence redesign using ProteinMPNN.^25^ We obtained three novel PET hydrolases, P4-a, P5-a, and P4-a-2, and were able to efficiently express P4-a-2 (also named as *Rs*PETase 1). Interestingly, *Rs*PETase 1 exhibits comparable PET hydrolytic activity to *Is*PETase and relatively high thermostability, demonstrating its great potential to be further engineered for industrial applications. These designer enzymes show low sequence and structural similarities compared to currently known PET hydrolases, manifesting the ability of deep learning algorithms to construct new-to-nature enzymes without relying on the existing protein scaffolds evolved by nature. This work paves an avenue to computationally build up the diversity of enzymes with known catalytic mechanisms, through which one can screen more robust and efficient enzymes (or proteins of interest) of industrial or pharmaceutical importance.

The successful replication of enzymatic activity by recapitulation of functional motifs with the same coordinates of native enzymes reaffirms that catalysis is accomplished by a small fraction of residues.^41, 42^ On the other hand, different protein scaffolds possess distinct inherent protein dynamics, including the atomic local fluctuations and large structure collective motions, which also significantly regulates the intrinsic activity of enzymes.^43^ The computational construction of isoenzymes with artificial protein architectures can yield isoenzymes with distinct protein dynamics that are absent in nature. With these isoenzymes we are able to pursue quantitative and mechanistic insights into the relationship between catalysis and protein dynamics.

The rapid development of the *de novo* design of proteins through deep learning models has also stoked the ambition to create small proteins or miniproteins with desired functions^15, 44, 45^. Small proteins have considerable advantages over larger ones: their genes are easier to synthesize, they take up fewer cell resources to express, they can be engineered to be highly stable, and they are potentially able to penetrate into tissues and cells.^46-48^ In the context of catalysis, smaller enzymes duplicate the desired catalytic activity using just the necessary residues and may benefit the cost-effectiveness of industrial applications. In this study, we intentionally generate new designs with shorter amino acid sequences compared to the original enzyme. The obtained active enzymes are 177 and 185 amino acids long, making them 30% smaller than the original enzyme LCC. The expression problems of some designs can be partly attributed to the smaller protein size, as the shortening of the polypeptide chains makes it more difficult to effectively bury hydrophobic amino acids in the protein core. The resulting exposure of hydrophobic areas increases the propensity for protein misfolding and aggregation.^45^ In addition, we found that the RF_joint_ model tends to “inpaint” missing regions with consecutively repeated and hydrophobic amino acids, making the design of small proteins even more challenging. Using multiple deep learning models for orthogonal validation can help overcome this limitation and increases the success rate of designs, but specific models trained with a collection of abundant small proteins would provide more accurate solutions. We anticipate that in the near future, the expansion of the small proteins database^49^ and the advancement of protein design approaches will boost our ability to create artificial small proteins and enzymes with greater accuracy and success rate.

## Methods

### Computational section

#### *In silico* modeling and analysis

The 3D structures of designed proteins were predicted by using ColabFold based on protein sequences. Molecular docking of 2-HE(MHET)_3_ to PET hydrolases was performed using AutoDockTools 1.5.6. The energetically favorable orientations of ligands binding to the targeted sites of enzymes were extracted and analyzed, and the docking conformations were then visualized and analyzed by Pymol 2.5.4.

#### MD simulations

MD simulations and analysis were performed using the GROMACS 2018 simulation package^50^ with the CHARMM36^51^ force field. The designed proteins were solvated with 0.15 M NaCl solution in a cubic box with a minimal distance of 1.0 nm from the box edge to the protein. The water molecules are modeled with the TIP3P model. To avoid unfavorable interactions, energy minimization was done using the steepest descent method prior to MD simulations. After equilibration for 100 ps in NVT ensemble and 100 ps in NPT ensemble at 300 K, a production run of 18 ns was conducted at 1 bar and 300 K. All MD simulations were conducted with a time step of 2 fs. The coordinates, energy, and velocity were stored every 0.5 ns for trajectory analysis. RMSD (including the motif RMSD and time-dependent RMSD), SASA (solvent accessible surface area), and R_g_ (radius of gyration) were calculated with tools in Gromacs. Simulation trajectories were visualized and analyzed using Pymol 2.5.4 and VMD 1.9.3.

#### Inpainting and refining the new sequences to scaffold the functional motifs

After extracting the functional motifs from LCC, the missing sequences were completed by the RF_joint_-guided inpainting approach.^14^ The retained amino acid sequences and structural information were used as input. During the inpainting, typically 10 repeated cycles were applied to generate high-quality inpainting sequences and structures. To further refine the inpainting results, we identified the problematic sequences (such as the SAAR and CHAA sequences) and large hydrophobic patches, and then deleted these regions. We utilized RF_joint_ to inpaint new sequences with the same length until there were no such flaws. Multiple rounds of inpainting were required to stepwise refine the sequences and structures.

#### Redesign of the protein sequence by ProteinMPNN

We used ProteinMPNN to redesign the whole sequences of the enzymes that exhibited PET hydrolytic activity.^25^ The open source code of the ProteinMPNN algorithm is available on github. The full protein backbone coordinates suggested by ColabFold were used as the input for ProteinMPNN to generate the new sequences. We generated 10 sequences for each target backbone (sampling temperature 0.1), and the protein structures of these sequences were predicted by ColabFold.

### Experimental section

#### Enzymatic assays

The enzyme activity was assayed based on the concentrations of soluble hydrolysates released from hydrolytic reactions. When using BHET as the substrate, a certain amount of purified mixed proteins was added to 1 mL phosphate buffer (50 mM, pH 8.0) containing 1 mg/mL BHET to initiate the reaction. After reaction at 25 °C, 800 rpm for 24 h in an incubator, the hydrolysates were analyzed by HPLC (see below). When using low-crystallinity PET powder as the substrate, enzymatic reactions were carried out with 6.0 μg/mL purified mixed proteins in the presence of 5 mg/ml of PET powder in phosphate buffer (50 mM, pH 8.0) at a given temperature under agitation at 800 rpm for 24 h. The reaction was quenched by heating to 100 °C, followed by the addition of the same volume of 1‰ aqueous trifluoroacetic acid. The reaction mixture was then filtered with polyethersulfone syringe filters (0.2 μm in pore size) to remove the remained PET powders. The filtrate was analyzed using an HPLC system equipped with an Eclipse Plus C18 column. The mobile phase was composed of methanol and 1‰ aqueous trifluoroacetic acid, and the flow rate was 1.0 mL/min. The hydrolysates were monitored by a diode-array detection at a wavelength of 240 nm, and the temperature of the column oven was 30 °C.

The Michaelis-Menten kinetics of P4-a-2 was assayed with 0.03 μM enzyme and various PET loads from 0–7.5 g/L at 50 °C in the shaking incubator at 800 rpm for 2 h. The concentrations of released hydrolysates were measured on a UV-Vis spectrophotometer at 240 nm, the products MHET, BHET, and TPA were regarded as the TPA equivalents (TPA_eq_), assuming having the same molar extinction coefficient of 17,000M^-1^cm^-1^.^52^ The *K*_m_ and *k*_*cat*_ values were determined by fitting the initial steady-state velocity to the Michaelis-Menten equation.

## Acknowledgement

This work was supported by the National Nature Science Foundation of China under grant number 32371325, and seed funding of China Petrochemical Corporation (Sinopec Group) under grant number 223260.

## Contributions

Y.Z. conceived and designed the research. Y.D. performed the computational design. Y.D. and S.Z. performed the experiments. Y.D and Y.Z. wrote the initial manuscript. Y.D., Y.Z., H.H, and X.K. discussed the results and revised the manuscript.

## Additional information

Supplementary information is available in the online version of the paper. Reprints and permissions information is available online at www.nature.com/reprints. Correspondence and requests for materials should be addressed to Y.Z.

## Supplementary Information

**PDF File**. Supplementary information including additional materials and methods, Supplementary Figs. 1–14, Supplementary Tables 1–6 and Supplementary References

## References

1. Katsimpouras, C. & Stephanopoulos, G. Enzymes in biotechnology: Critical platform technologies for bioprocess development. Curr. Opin. Biotechnol. 69, 91–102 (2021).

2. Bell, E.L., et al. Biocatalysis. Nat. Rev. Methods Primers 1, 46 (2021).

3. Buller, R., et al. From nature to industry: Harnessing enzymes for biocatalysis. Science 382, (2023).

4. Riziotis, I. G., Ribeiro, A. J. M., Borkakoti, N. & Thornton, J. M. Conformational variation in enzyme catalysis: a structural study on catalytic residues. J. Mol. Biol. 434, 167517 (2022).

5. Gherardini, P. F., Wass, M. N., Helmer-Citterich, M. & Sternberg, M. J. Convergent evolution of enzyme active sites is not a rare phenomenon. J. Mol. Biol. 372, 817–845 (2007)

6. Jiang, L., et al. De novo computational design of retro-aldol enzymes. Science 319, 1387–1391 (2008).

7. Röthlisberger, D., et al. Kemp elimination catalysts by computational enzyme design. Nature 453, 190–195 (2008).

8. Robles-Martín, A., et al. Sub-micro- and nano-sized polyethylene terephthalate deconstruction with engineered protein nanopores. Nat. Catal. 6, 1174–1185 (2023).

9. Kalvet, I., et al. Design of heme enzymes with a tunable substrate binding pocket adjacent to an open metal coordination site. J. Am. Chem. Soc. 145, 14307–14315 (2023).

10. Alexander, P. A., et al. A minimal sequence code for switching protein structure and function. Proc. Natl. Acad. Sci. USA. 106, 21149–21154 (2009).

11. Pan, X. & Kortemme, T. Recent advances in de novo protein design: Principles, methods, and applications. J. Biol. Chem. 296, 100558 (2021).

12. Senior, A. W., et al. Improved protein structure prediction using potentials from deep learning. Nature 577, 706–710 (2020).

13. Jumper, J., et al. Highly accurate protein structure prediction with AlphaFold. Nature 596, 583–589 (2021).

14. Wang, J., et al. Scaffolding protein functional sites using deep learning. Science 377, 387–394 (2022).

15. Yeh, A. H., et al. De novo design of luciferases using deep learning. Nature 614, 774–780 (2023).

16. Madani, A., et al. ProGen: Language modeling for protein generation. bioRxiv preprint, doi: 10.1101/2020.03.07.982272.

17. Robinson, S. L., Piel, J. & Sunagawa, S. A roadmap for metagenomic enzyme discovery. Nat. Prod. Rep. 38, 1994–2023 (2021).

18. Ferrer, M., et al. Estimating the success of enzyme bioprospecting through metagenomics: current status and future trends. Microb. Biotechnol. 9, 22–34 (2016).

19. Eiamthong, B., et al. Discovery and genetic code expansion of a polyethylene terephthalate (PET) hydrolase from the human saliva metagenome for the degradation and Bio-Functionalization of PET. Angew Chem. Int. Ed. 61:e202203061 (2022).

20. Yoshida, S., et al. A bacterium that degrades and assimilates poly(ethylene terephthalate). Science 351, 1196–1199 (2016).

21. Buchholz, P. C. F., et al. Plastics degradation by hydrolytic enzymes: The plastics-active enzymes database-PAZy. Proteins 90, 1443–1456 (2022).

22. Joo, S., et al. Structural insight into molecular mechanism of poly(ethylene terephthalate) degradation. Nat. Commun. 9, 382 (2018).

23. Sulaiman, S., et al. Isolation of a novel cutinase homolog with polyethylene terephthalate-degrading activity from leaf-branch compost by using a metagenomic approach. Appl. Environ. Microbiol. 78, 1556–1562 (2012).

24. Mirdita, M., et al. ColabFold: making protein folding accessible to all. Nat. Methods 19, 679–682 (2022).

25. Dauparas, J., et al. Robust deep learning-based protein sequence design using ProteinMPNN. Science 378, 49–56 (2022).

26. Sulaiman S., et al. Crystal structure and thermodynamic and kinetic stability of metagenome-derived LC-cutinase. Biochemistry 53, 1858–1869 (2014).

27. Han X., et al. Structural insight into catalytic mechanism of PET hydrolase. Nat. Commun. 8, 2106 (2017).

28. Riziotis, I. G., et al. The 3D modules of enzyme catalysis: Deconstructing active sites into distinct functional Entities. J. Mol. Biol. 435, 168254 (2023).

29. Jerves, C., et al. Reaction mechanism of the PET degrading enzyme PETase studied with DFT/MM molecular dynamics simulations. ACS Catal. 11, 11626–11638 (2021).

30. Madani, A., et al. Structure-informed language models are protein designers. bioRxiv preprint, doi: https://arxiv.org/abs/2302.01649

31. Zheng, Z., et al. Understanding the role of hydrophobic patches in protein disaggregation. Phys Chem Chem Phys 23, 12620–12629 (2021).

32. Mahmoud, S. A. & Chien, P. Regulated proteolysis in bacteria. Annu. Rev. Biochem. 87, 677–696 (2018).

33. Schlegel, S., et al. Optimizing membrane protein overexpression in the Escherichia coli strain Lemo21(DE3). J. Mol. Biol. 423, 648–659 (2012).

34. Butt, T. R., et al. SUMO fusion technology for difficult-to-express proteins. Protein Expr. Purif. 43, 1–9 (2005).

35. Baker, D. What has de novo protein design taught us about protein folding and biophysics? Protein Sci. 28, 678–683 (2019).

36. Baath, J. A., et al. Comparative biochemistry of four Polyester (PET) hydrolases. ChemBioChem 22, 1627–1637 (2021).

37. Bell, E. L., et al. Directed evolution of an efficient and thermostable PET depolymerase. Nat. Catal. 5, 673–681 (2022).

38. Lu, H., et al. Machine learning-aided engineering of hydrolases for PET depolymerization. Nature 604, 662–667 (2022).

39. Arnal, G., et al. Assessment of four engineered PET degrading enzymes considering large-scale industrial applications. ACS Catal. 13, 13156–13166 (2023).

40. Kaabel, S., et al. Enzymatic depolymerization of highly crystalline polyethylene terephthalate enabled in moist-solid reaction mixtures. Proc. Natl. Acad. Sci. USA 118 (2021).

41. Ribeiro, A. J. M., et al. A global analysis of function and conservation of catalytic residues in enzymes. J. Biol. Chem. 295, 314–324 (2020).

42. Bartlett, G. J., et al. Analysis of catalytic residues in enzyme active sites. J. Mol. Biol. 324, 105–121 (2002).

43. Nam, K. & Wolf-Watz, M. Protein dynamics: The future is bright and complicated! Struct. Dyn. 10, 014301 (2023).

44. Ozga, K. & Berlicki, L. Design and engineering of miniproteins. ACS Bio. Med. Chem. Au 2, 316–327 (2022).

45. Kim, D. E., et al. De novo design of small beta barrel proteins. Proc. Natl. Acad. Sci. USA 120, e2207974120 (2023).

46. Jiang, M., Lou, H. & Hou, W. Microproteins: from behind the scenes to the spotlight. Genome Instability & Disease 2, 225–239 (2021).

47. Crook, Z. R., Nairn, N. W. & Olson, J. M. Miniproteins as a powerful modality in drug development. Trends Biochem. Sci. 45, 332–346 (2020).

48. Cao, L., et al. De novo design of picomolar SARS-CoV-2 miniprotein inhibitors. Science 370, 426–431 (2020).

49. Li, Y., et al. SmProt: A reliable repository with comprehensive annotation of small proteins identified from ribosome profiling. Genomics. Proteomics Bioinformatics 19, 602–610 (2021).

50. Lemkul, J. A. From proteins to perturbed hamiltonians: A Suite of tutorials for the GROMACS-2018 molecular simulation package [Article v1.0]. Living Journal of Computational Molecular Science (2019).

51. Huang, J., et al. CHARMM36: An improved force field for folded and intrinsically disordered proteins. Biophys. J. 112, 175a–176a (2017).

52. Aristizábal-Lanza, L., et al. Comparison of the enzymatic depolymerization of polyethylene terephthalate and AkestraTM using Humicola insolens cutinase. Front. Chem. Eng. 4, 1048744 (2022).

