## supplementary information for "Replicating enzymatic activity by positioning active sites with synthetic protein scaffolds"

#### Supplementary Materials and Methods

**Materials.** Bis(2-Hydroxyethyl) terephthalate (BHET) was purchased from Shanghai Macklin Biochemical Technology Co., Ltd. The low-crystallinity PET (LC-PET) was obtained from post-consumer Coca-Cola bottles. The shoulder part of the bottle was cut into 2 × 5 mm pieces, and then microsized into 50–150 μm powders by ball milling. Kanamycin and β-D-1-thiogalactopyranoside (IPTG) were purchased from Yuanye Bio-Technology Co., Ltd. (Shanghai, China). Luria−Bertani (LB) was purchased from Sangon Biotech Co., Ltd. (Shanghai, China). All chemicals and reagents are of analytical grade.

All DNA primers and gene synthesis were completed by the Beijing Genomics Institute. DNA modifying enzymes and T4 DNA ligase were purchased from New England Biolabs. Plasmid extraction and gel purification kits were bought from Omega. Lemo21 (DE3) competent *E. coli* cells were purchased from Beijing Zoman Biotechnology Co., Ltd. The *E. coli* BL21 (DE3), *E. coli* BL21 (DE3) PLysS cells, *E. coli* Trans5α cells, plasmid pET28a (+) and DNA polymerase (2× PrimeSTAR Max Premix) were purchased from Tsingke Biology Co., Ltd. Cells were cultured in Luria–Bertani (LB) medium.

**Methods**

**Plasmid construction and enzyme expression.** All designs tested in *E. coli* were cloned, expressed and purified using standard methods. All genes encoding the designer proteins were synthesized by Beijing Genomics Institute. Genes encoding the first-round designed proteins (P1–P10) and the refined proteins (P4-a, P5-a and P7-a) were inserted into modified pET28a (+) vectors containing a N-terminal $His_6$-tags. Genes encoding proteins P4-a and P5-a were also inserted into modified pET28a (+) vectors containing N-terminal SUMO cleavage sites and a $His_6$-tags. Genes encoding proteins P4-a-1, P4-a-2, P5-a-1, and P5-a-2 were inserted into pET29b (+) vectors. The constructed plasmids were transformed into corresponding *E. coli* stains (Supplementary Table 5). After an overnight culture on a Luria-Bertani (LB) agar plate containing 50 μg/mL kanamycin, a single colony was selected and inoculated into 3 mL of LB medium with 50 μg/mL kanamycin and incubated overnight at 37 °C with shaking at 200 rpm. The cells were then cultivated in 300 mL LB medium containing 50 μg/mL kanamycin at 37 °C/200 rpm until an $OD_{600}$ reached above 0.6–0.8. Enzymes expression was induced by 0.5 mM IPTG at 16 °C for 18 h. The bacterial cells were harvested by centrifugation at 4000 × g for 30 min at 4 °C and resuspended in a lysis buffer (50 mM Tris-HCl, 300 mM NaCl and 10 mM imidazole at pH 8.0). The resuspended cells are disrupted by a benchtop high pressure homogenizer at 4 °C. Cell debris was removed by centrifugation at 4 °C, 10000 × g for 30 min. The supernatant was incubated with a Ni-NTA resin for 1 h at 4 °C. The non-specifically adsorbed proteins were removed by washing with 50 mL wash buffer (50 mM Tris-HCl, 300 mM NaCl and 50 mM imidazole at pH 8.0). Then, the His-tagged protein was eluted with elution buffer (50 mM Tris-HCl, 300 mM NaCl and 300 mM imidazole at pH 8.0). The purified proteins were concentrated using a 10 kDa Amicon Ultra centrifuge tube (Millipore, Burlington, USA) and was then exchanged with sodium phosphate buffer (50 mM, pH 8.0) by passing through the HiTrap desalting column. Before the enzyme reaction, the enzyme concentration was quantified by the BCA Protein Assay Kit.

**Cloning:** P4-a and P5-A fused to SUMO were cloned into pET28a/His-SUMO A vector. The genes gP4-a /gP5-a for P4-a and P5-a were amplified from plasmids synthesized by the Beijing Genomics Institute. All primer informations are listed in Supplementary Table 6**.** To obtain linearized pET28a His-SUMO A vector, a 50 μL polymerase chain reaction (PCR) system was prepared containing 9.5 ng pET28a/His-SUMO A vector plasmid, 0.5 μM primers (for both forward and reverse primers) and 25 μL 2× High-Fidelity Master Mix. The PCR was carried out on a Thermo Scientific Arktik thermal cycler (98 ℃ for 2 min; 98 ℃ for 10 s, 55 ℃ for 5 s, 72 ℃ for 30 s, 33 cycles; 72 ℃ for 5 min; 4 ℃ hold). The PCR product was digested by BamH I and XhoI (100 U, NEB) at 37 ℃ for 2 h to degrade the template vector, and was then purified using PCR Purification Kit.

The linearized pET28a/His-SUMO A vector and insert genes (gP4-a/gP5-a) were assembled at 22 ℃ for 1.5 h in a 10 μL reaction system, containing the vector (5 ng), inserts (1.5 ng gP4-a or 1.5 ng gP5-a) and 1 μL T4 DNA ligase with 1 μL 10× T4 DNA ligase buffer. The assembly products were transformed into 25 μL *E. coli* Trans5α Cells by electroporation using Eppendorf Eporator. The cells were then mixed with 400 μL LB medium and incubated at 37 ℃ with shaking for 50 min. The cells were cultured on LB agar plates with ampicillin (0.05 mg/mL), and the colonies were selected were inoculated in 5 mL LB broth with antibiotics and incubated at 37 ℃ with 200 rpm shaking overnight. The plasmid DNA was extracted using Plasmid Mini Kit. Colonies resulting from transformation of the ligation were screened by colony PCR, restriction enzyme digestion and all hits were sequenced for complete verification. The primers for DNA sequencing were universal primers or designed based on the sequences of P4-a/P5-a (see Supplementary Table 6).

**Size exclusion chromatography (SEC).** AKTA pure M with UNICORN 6.3.2 Workstation control (GE Healthcare) coupled with a Superdex 75 Increase 10/300 GL column and buffer (10 mM Tris, 500 mM NaCl, 5 mM DTT) was used for size exclusion chromatography.

**Differential scanning fluorescence (DSF) analysis Tm values of Protein.** The protein sample solution was filled into 3 nanoDSF Grade Standard Capillaries (NanoTemper Technologies) and integrated in a Prometheus NT.48 device (Nanotemper Technologies) controlled by the PR. ThermControl software (version 2.1.2). Excitation power was pre-adjusted to get fluorescence readings above 2000 RFU for F330 and F350, and samples were heated from 40 ℃ to 80 ℃ with a slope of 0.5 ℃/min. An XLSX file with "processed data" was exported from the PR. ThermControl software and used for further analysis.

### Supplementary Figures

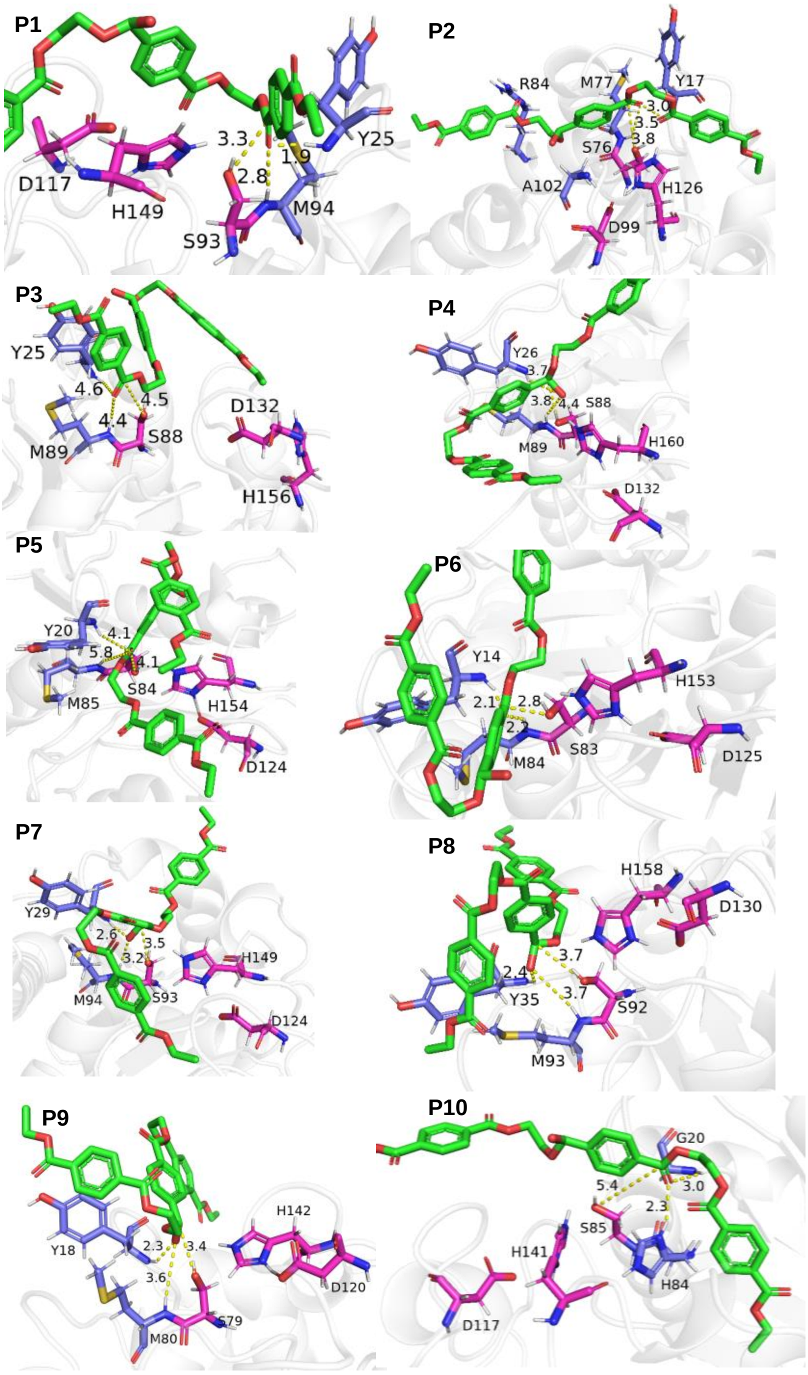


**Supplementary Figure 1.** Representative molecular docking of 2-HE(MHET)$_3$ (green stick model) with the virtual enzymes P1 to P10. The catalytic triad, Ser, His, and Asp are highlighted in magenta. The residues which can potentially form the oxyanion hole are colored in blue-grey. The distance between the oxygen atom from the side chain of the Ser residue and the carbonyl carbon of the ester bond is indicated in each docking with dashed lines. Oxygen atoms are colored in red, hydrogen atoms in white, nitrogen atom in blue, and sulphur atoms in yellow.

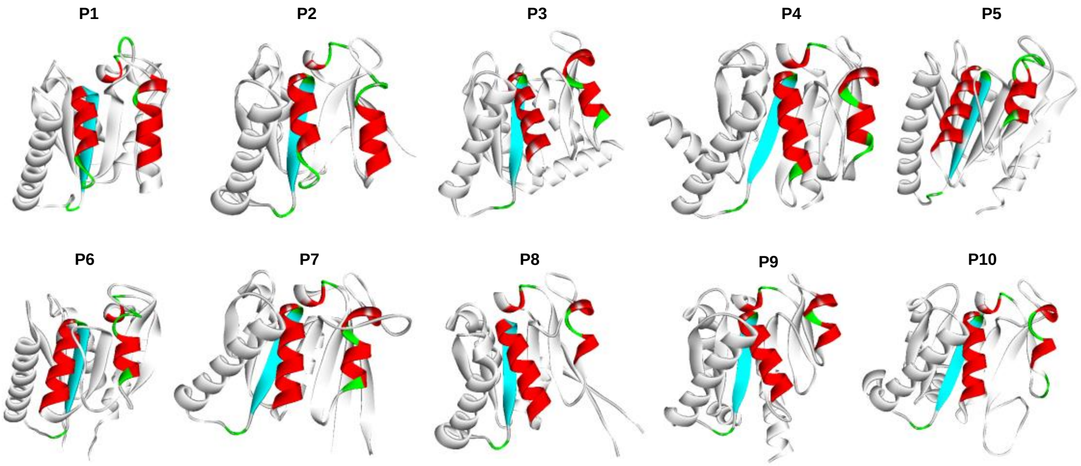


**Supplementary Figure 2.** The predicted 3D structure of the 10 virtual enzymes generated in the first round of design. The newly generated protein scaffolds are colored in gray, while the retained regions from the template enzyme LCC are colored in red, green and blue.

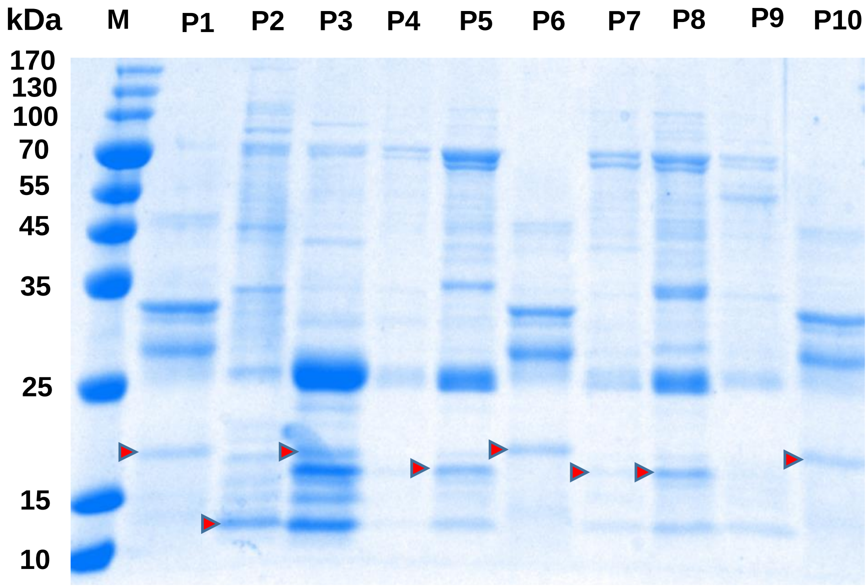


**Supplementary Figure 3.** SDS-PAGE analysis of the expressed P1–P10 in *E. Coli* cells after purification by Ni-NTA chromatography. The expected bands of target proteins are indicated with the red arrows. Although the bands of P1, P2, P3, P5, P6, P7, P8, P10 are shown in the gel, their expression levels are too low to be further purified.

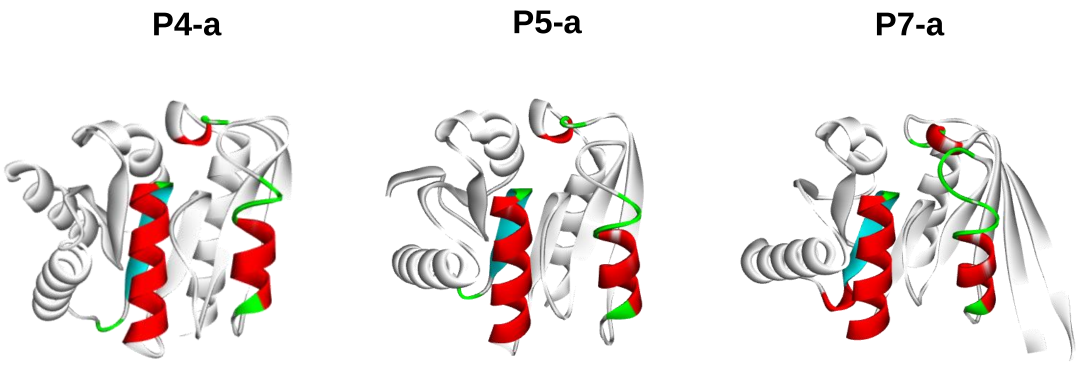


**Supplementary Figure 4.** The 3D structure of the redesigned enzymes using iterative $RF_{joint}$. The regenerated protein scaffolds are colored in grey while the retained regions from the template enzyme LCC are colored in red, green and blue.

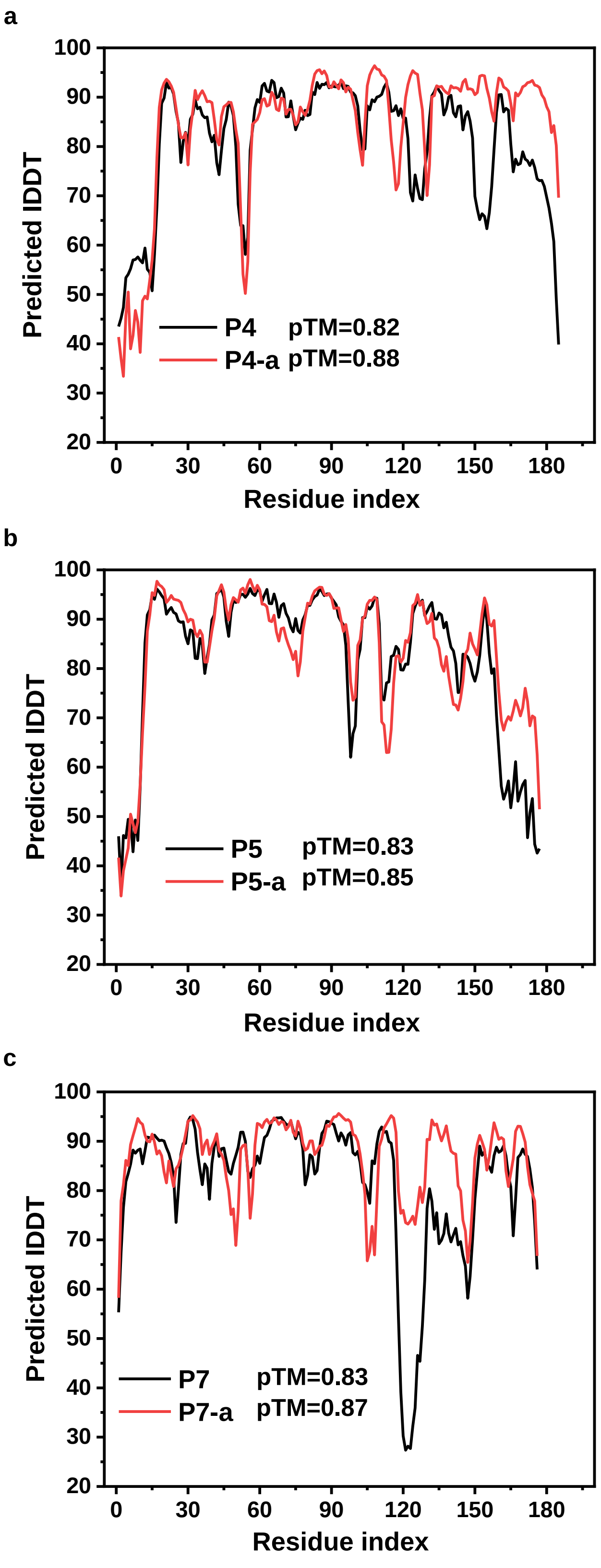


**Supplementary Figure 5.** Comparisons of the pLDDT values and pTM-scores before and after sequence refinement by $RF_{joint}$ inpainting. a, P4 (black curve) and P4-a (red curve); b, P5 (black curve) and P5-a (red curve); c, P7 (black curve) and P7-a (red curve).

**a**

```
LCC                1 SNPYQRGPNPTRSALTADGPFSVATYTVSRLSVSGFGGGVIYYPTGTSLT     50
                     |       |..|:...|:              |...|.|.:
P4-a               1 S-------NAERARALAE--------------VRARGRGAL---------     20

LCC               51 FGGIAMSPGYTADASSLAWLGRRLASHGFVVLVINTNSRFDYPDSRASQL    100
                         .:..||||||||||||||||||. |.||::|..    |||||||||
P4-a              21 ----VLDGGYTADASSLAWLGRRLASL-FGVLLVNAR----YPDSRASQL     61

LCC              101 SAALNYLRTSSPSAVRARLDANRLAVAGHSMGGGGTLRIAEQNPSLKAAV    150
                     |||||||||...|   ..|||||||||||||||||||||||||......:
P4-a              62 SAALNYLRTQLAS---LGLDANRLAVAGHSMGGGGTLRIAEQNRGRPRVL    108

LCC              151 PLTPWHTDKTFNTSVPVLIVGAEADTVAPVSQHAIPFYQNLPSTTPKVYV    200
                     ..|||.:.........|| .|..:|||||||||||||||||.....:|..
P4-a             109 AFTPWESGPARGGRADVL-AGRGSDTVAPVSQHAIPFYQNLRGGRLRVLP    157

LCC              201 ELDNASHFAPNSNNAAISVYTISWMKLWVDNDTRYRQFLCNVNDPALSDF    250
                         |.|||||..:.|:.....:.::.|...
P4-a             158 ----APHFAPNRGSPAVVAAARAALEAWEAQ-------------------    184

LCC              251 RTNNRHCQ    258
                            |
P4-a             185 -------Q    185
```

**b**

```
LCC                1 SNPYQRGPNPTRSALTADGPFSVATYTVSRLSVSGFGGGVIYYPTGTSLT     50
                     ||.         ..|.|....::..|.
P5-a               1 SNA---------QLLRAGAQLALVLYA-----------------------     18

LCC               51 FGGIAMSPGYTADASSLAWLGRRLASHGFVVLVINTNSRFDYPDSRASQL    100
                             ||||||||||||||||||.|..||:::...   |||||||||
P5-a              19 --------GYTADASSLAWLGRRLASLGLAVLLLSGGG---YPDSRASQL     57

LCC              101 SAALNYLRTSSPSAVRARLDANRLAVAGHSMGGGGTLRIAEQNPSLKAAV    150
                     |||||||||...|   ..|||||||||||||||||||||||||.....|:
P5-a              58 SAALNYLRTQLAS---LGLDANRLAVAGHSMGGGGTLRIAEQNRGGLLAL    104

LCC              151 PLTPWHTDKTFNTSVPVLIVGAEADTVAPVSQHAIPFYQNLPSTTPKVYV    200
                     ..|||...:....   ||::|  :||||||||||||||||......::..
P5-a             105 AFTPWEGGRAGRA---VLVLG--SDTVAPVSQHAIPFYQNARPGGFRLLR    149

LCC              201 ELDNASHFAPNSNNAAISVYTISWMKLWVDNDTRYRQFLCNVNDPALSDF    250
                     ...|  |||||:..|.                            .||.:.
P5-a             150 VPGN--HFAPNNGRAL----------------------------DALEEL    169

LCC              251 RTNNRHCQ    258
                     ....:..|
P5-a             170 LQLLQQQQ    177
```

**c**

```
LCC                1 SNPYQRGPNPTRSALTADGPFSVATYTVSRLSVSGFGGGVIYYPTGTSLT     50
                     |       |.:......|....:|.:....|.:..
P7-a               1 S-------NCSVEVRRLDSLEELADFLEEELGLDD---------------     28

LCC               51 FGGIAMSPGYTADASSLAWLGRRLASHGFVVLVINTNSRFDYPDSRASQL    100
                        :|::.||||||||||||||||||     |::|.|.   |||||||||
P7-a              29 ---VALASGYTADASSLAWLGRRLAS-----LLVNGNG---YPDSRASQL     67

LCC              101 SAALNYLRTSSPSAVRARLDANRLAVAGHSMGGGGTLRIAEQNPSLKAAV    150
                     |||||||||:.|.     ||||||||||||||||||||||||| .|..|:
P7-a              68 SAALNYLRTNRPG-----LDANRLAVAGHSMGGGGTLRIAEQN-GLPGAL    111

LCC              151 PLTPWHTDKTFNTSVPVLIVGAEADTVAPVSQHAIPFYQNLPSTTPKVYV    200
                     ..|||..    .....:|::|  |||||||||||||||||||...    :
P7-a             112 VFTPWGP----GGGGRLLLLG--ADTVAPVSQHAIPFYQNLPGGR----L    151

LCC              201 ELDNASHFAPNSNNAAISVYTISWMKLWVDNDTRYRQFLCNVNDPALSDF    250
                     .|....|||||:....:.:..::
P7-a             152 ALSGDGHFAPNNGRGRVLLLEVT---------------------------    174

LCC              251 RTNNRHCQ    258
                           ||
P7-a             175 ------CQ    176
```

**Supplementary Figure 6.** Pairwise sequence alignment of the newly designed enzymes with respect to the template enzyme LCC using the online tool provided by EMBL-EBI.[1] The sequence identity of P4-a, P5-a and P7-a with LCC is 41%, 40% and 40%, respectively. The sequence similarity of P4-a, P5-a and P7-a with LCC is 47%, 46% and 47%, respectively.

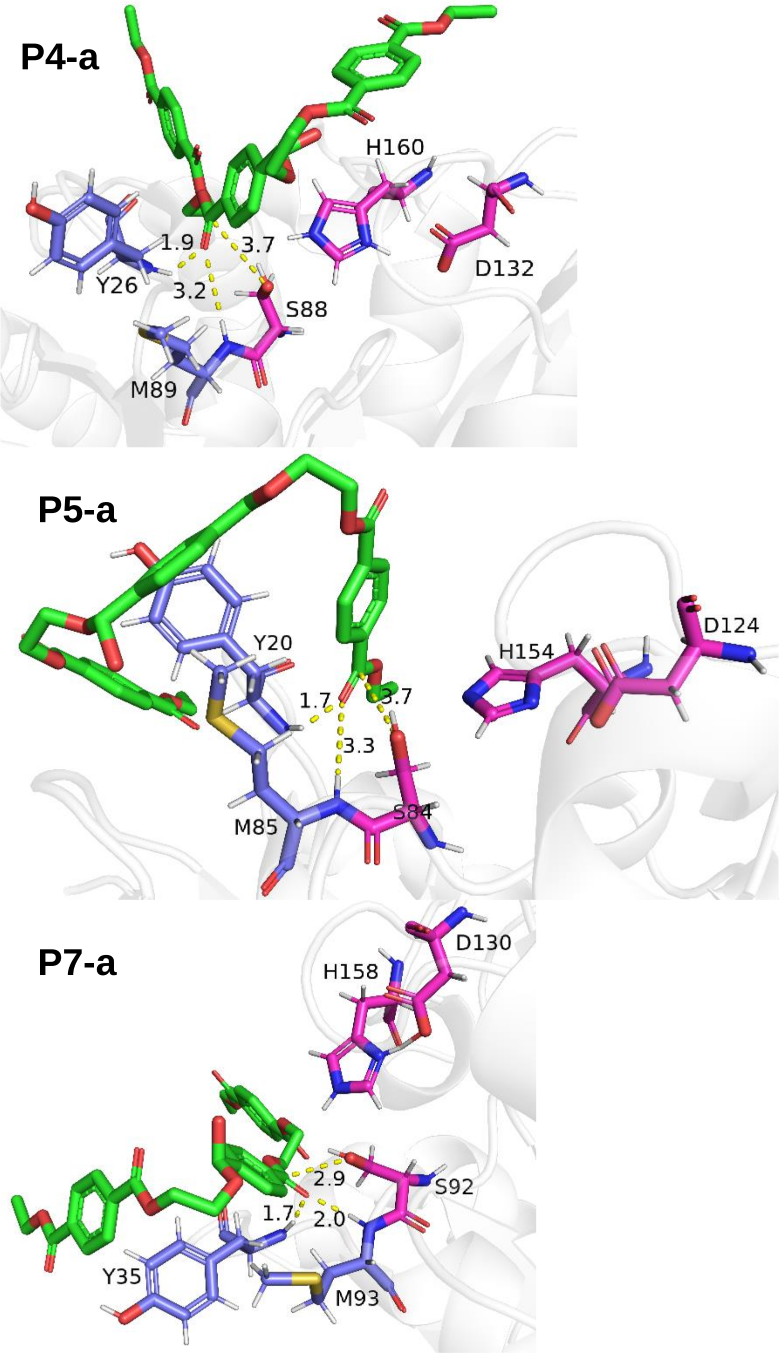


**Supplementary Figure 7.** Representative molecular docking of 2-HE(MHET)$_3$ (green colored stick model) with the virtual enzymes P4-a, P5-a and P7-a. The catalytic residues (Ser-His-Asp) are highlighted in magenta. The residues which can potentially form the oxyanion hole are colored in blue-grey. The distance between the oxygen atom from the side chain of the Ser residue and the carbonyl carbon of the ester bond is indicated in each docking with dashed lines. Oxygen atoms are colored in red, hydrogen atoms in white, nitrogen atom in blue, and sulfur atoms in yellow.

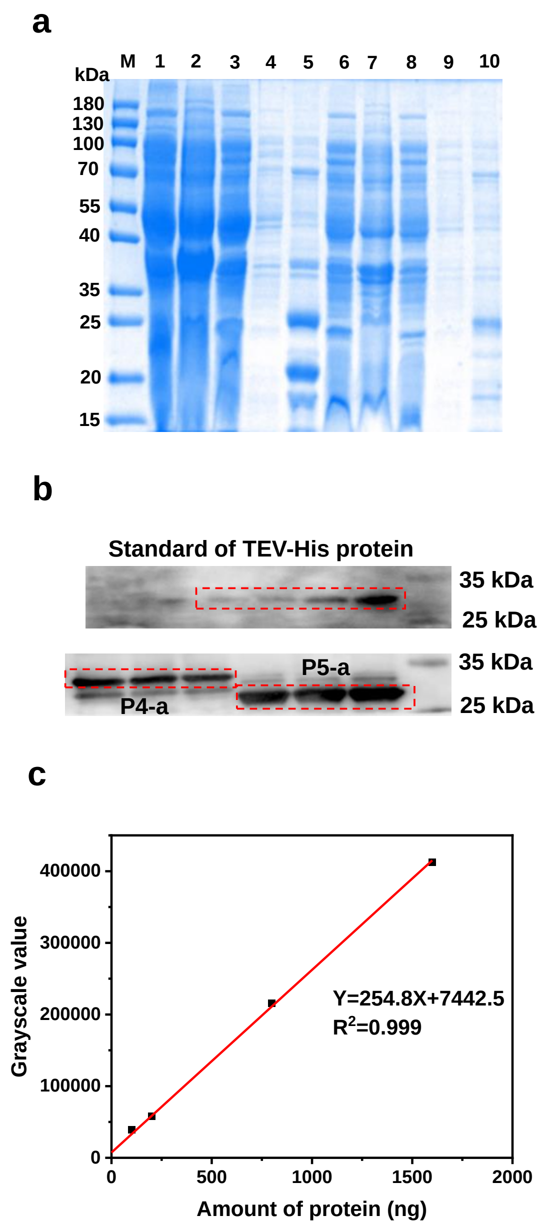


**Supplementary Figure 8. a,** SDS-PAGE analysis of the expression of His-tagged P4-a and P5-a in *E. coli* cells. Lane 1-5 for the analysis of P4-a: soluble fraction of the cell lysate (lane 1); precipitates of the cell lysate (lane 2); flow-through fraction (lane 3); elutes by 50 mM imidazole (lane 4) and by 300 mM imidazole (lane 5) from Ni-NTA affinity chromatography. Lane 6-10 for the analysis of P5-a: soluble fraction of the cell lysate (lane 6); precipitates of the cell lysate (lane 7); flow-through fraction (lane 8); elutes by 50 mM imidazole (lane 9) and by 150 mM imidazole (lane 10) from Ni-NTA affinity chromatography. **b,** Western blotting analysis of the His-tagged P4-a, P5-a. TEV-His protein was used as the protein standard for concentration assay. The proteins were visualized by anti-His6 western blotting. **c,** Standard curve (grey value as a function of protein amount) calibrated with TEV protein.

**a**

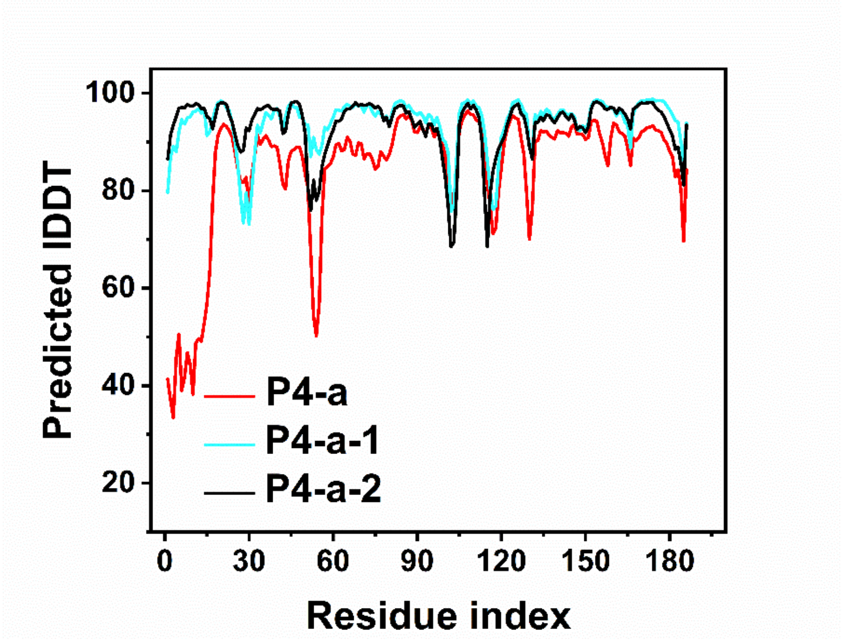


**b**

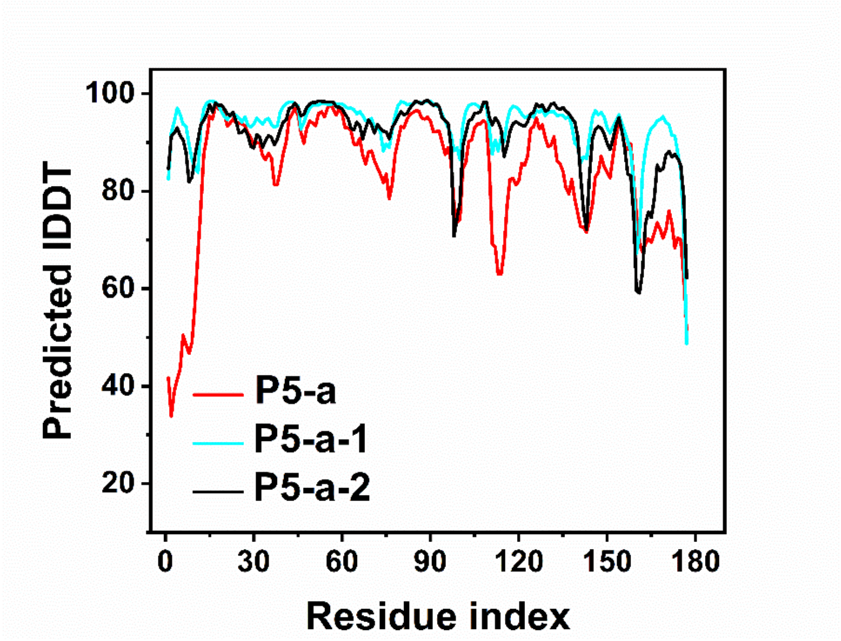


**Supplementary Figure 9.** Comparisons of the pLDDT values before and after ProteinMPNN rescue. a, P4-a (red curve), P4-a-1 (blue curve) and P4-a-2 (black curve); b, P5-a (red curve), P5-a-1 (blue curve) and P5-a-2 (black curve).

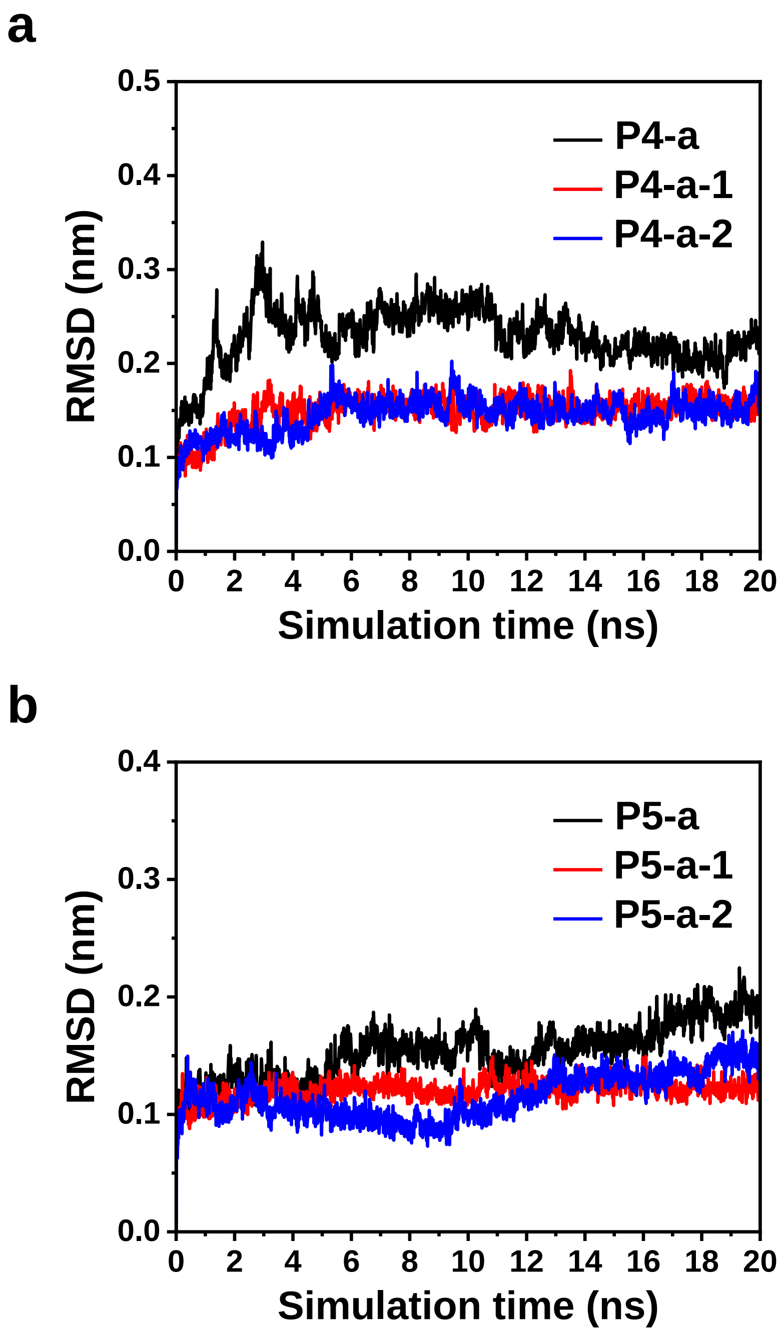


**Supplementary Figure 10.** Time-course RMSD fluctuations of **a**, P4-a, P4-a-1 and P4-a-2; and **b**, P5-a, P5-a-1 and P5-a-2.

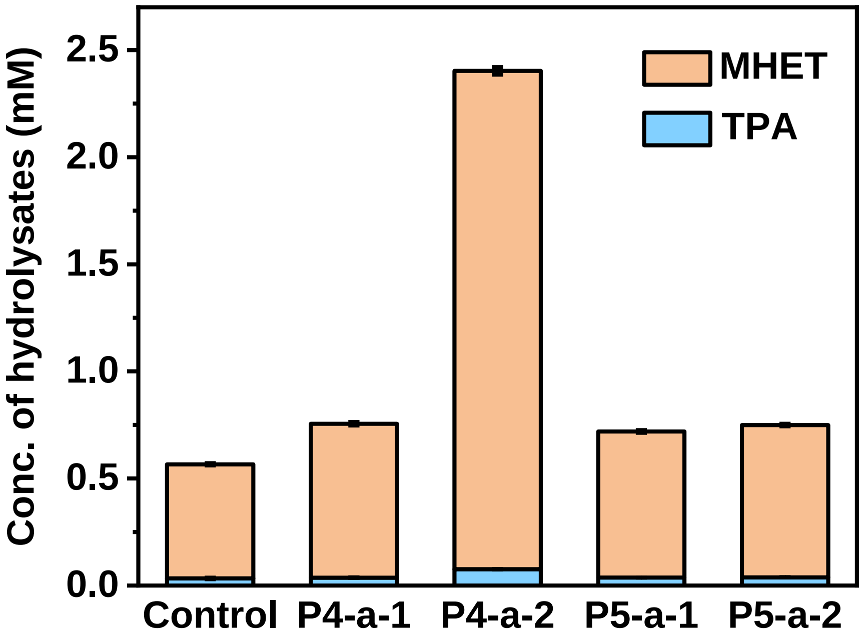


**Supplementary Figure 11.** The hydrolysis of BHET by the ProteinMPNN rescued enzymes (assayed with the eluted fractions from the Ni-NTA affinity column). Conditions: 6.0 μg/mL eluted proteins, 1.0 mg/mL BHET, 25 °C for 24 h. The hydrolytic products TPA and MHET were quantified by HPLC. Error bars represent the standard deviations of three measurements.

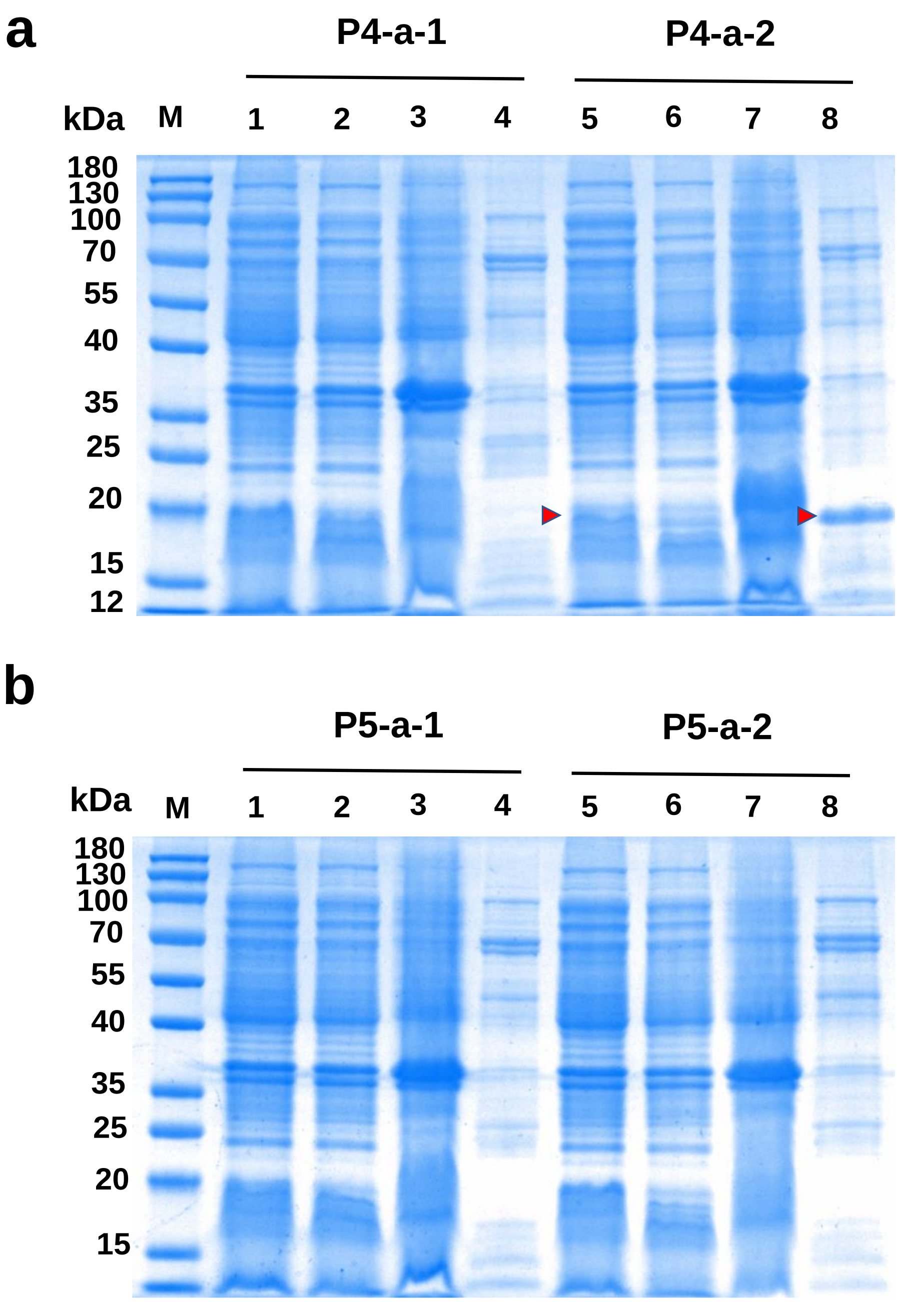


**Supplementary Figure 12.** SDS-PAGE plot of Coomassie brilliant blue stained numbered P4-a-1, P4-a-2, P5-a-1 and P5-a-2 proteases. **a** Lane 1: Soluble lysate of P4-a-1; Lane 2: precipitates of P4-a-1; Lane 3: flow-through fluid of P4-a-1; Lane 4: eluted proteins P4-a-1 from the Ni-NTA affinity chromatography; Lane 5: Soluble lysate of P4-a-2; Lane 6: precipitates of P4-a-2; Lane 7: flow-through fluid of P4-a-2; Lane 8: eluted proteins P4-a-2 from the Ni-NTA affinity chromatography; **b** Lane 1: Soluble lysate of P5-a-1; Lane 2: precipitates of P5-a-1; Lane 3: flow-through fluid of P5-a-1; Lane 4: eluted proteins P5-a-1 from the Ni-NTA affinity chromatography; Lane 5: Soluble lysate of P5-a-2; Lane 6: precipitates of P5-a-2; Lane 7: flow-through fluid of P5-a-2; Lane 8: eluted proteins P5-a-2 from the Ni-NTA affinity chromatography;

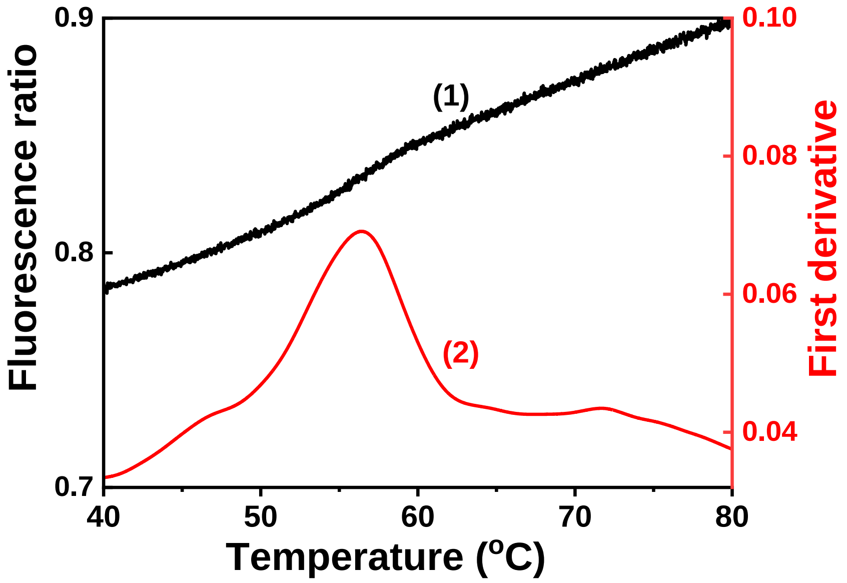


**Supplementary Figure 13. DSF scanning curves of P4-a-2.** The black curve represents the ratio of intrinsic fluorescence (350 nm: 330 nm) for DSF, and the red curve shows the first derivative.

**Supplementary Figure 14.** Pairwise sequence alignment of the P4-a-2 enzyme with respect to (a) LCC and (b) *Is*PETase. The sequence identity and similarity of P4-a-2 compared to LCC is 21% and 34%, respectively. The sequence identity and similarity of P4-a-2 compared to *Is*PETase is 18% and 25%, respectively. **(Sequences of P4-a-2 is not shown here; it can be provided upon request).**

#### Supplementary Tables

**Supplementary Table 1: Detailed information of the designer enzymes P1-P10, P4-a, P5-a and P7-a.**

The molecular weight, amino acid length, motif RMSD compared to the retained template structures, pLDTT and TM-score given by AlphaFold 2, the size and depth of the active pocket, and calculated binding energy with 2-HE(MHET)$_3$ are summarized below.

| Protein | Molecular Weight (kDa) | Length (aa) | Binding energy (kcal/mol) | Motif RMSD (Å) | pLDDT | Volume (Å$^3$) * | Depth (Å) * |
|---|---|---|---|---|---|---|---|
| LCC | 27.78 | 258 | -2.68 | - | - | 245 | 13.8 |
| P1 | 16.26 | 160 | -3.57 | 1.81 | 71.32 | 191 | 9.2 |
| P2 | 14.28 | 140 | -2.67 | 2.13 | 70.62 | 534 | 14.9 |
| P3 | 18.06 | 176 | -0.82 | 2.00 | 79.72 | 80 | 6.7 |
| P4 | 19.24 | 185 | -2.3 | 1.74 | 80.50 | 213 | 11.7 |
| P5 | 18.17 | 177 | -1.49 | 1.05 | 82.34 | 222 | 13.8 |
| P6 | 19.31 | 180 | -1.84 | 1.29 | 89.79 | 221 | 13.1 |
| P7 | 17.93 | 176 | -1.74 | 1.32 | 81.89 | 201 | 10.6 |
| P8 | 17.22 | 169 | -2.31 | 2.06 | 74.76 | 222 | 13.3 |
| P9 | 17.82 | 173 | -1.67 | 2.22 | 83.14 | 249 | 15.3 |
| P10 | 18.68 | 186 | -3.66 | 1.92 | 61.64 | 103 | 6.8 |
| P4-a | 19.43 | 185 | -2.74 | 1.22 | 84.39 | 215 | 11.8 |
| P5-a | 18.47 | 177 | -1.18 | 1.05 | 83.72 | 252 | 13.1 |
| P7-a | 18.34 | 176 | -2.57 | 1.11 | 86.92 | 221 | 12.6 |

* The size of active pocket and substrate binding energy were calculated by Protein plus.[2]

**Supplementary Table 2. Amino acid sequences of the wild-type LCC and designer enzymes.**

The underlined sequences are retained for computational rescaffolding, and the catalytic triad is highlighted in red.

**Wild-type LCC**

SNPYQRGPNPTRSALTADGPFSVATYTVSRLSVSGFGGGVIYYPTGTSLTFGGIAMSPGYTADASSLAWLGRRLASHGFVVLVINTNSRFDYPDSRASQLSAALNYLRTSSPSAVRARLDANRLAVAGH**S**MGGGGTLRIAEQNPSLKAAVPLTPWHTDKTFNTSVPVLIVGAEA**D**TVAPVSQHAIPFYQNLPSTTPKVYVELDNAS**H**FAPNSNNAAISVYTISWMKLWVDNDTRYRQFLCNVNDPALSDFRTNNRHCQ

**Amino acid sequences of designer enzymes**

**> P1**

SNTLGLGLELALLRGAGALALAGGYTADASSLAWLGRRLASELGLAVALVPAGGYPDSRASQLSAALNYLRTRLAARGLSRLDANRLAVAGH**S**MGGGGTLRIAEQGLLLALTPWGG**D**TVAPVSQHAIPFYQNLLRLGLRGVVLVVGGN**H**FAPNNGLGLCQ

**> P2**

SNSLDGLGDAFALAAGYTADASSLAWLGRRLASLLVPAGYPDSRASQLSAALNYLRTQLASELGLDANRLAVAGH**S**MGGGGTLRIAEQGLLLVLTPWG**D**TVAPVSQHAIPFYQNCGLPLLLAGPG**H**FAPNNGLGLGLNCQ

**> P3**

SNAALLLALLAGLRQARALALASGYTADASSLAWLGRRLASELGLGVLLVNGNGYPDSRASQLSAALNYLRTNRPGLDANRLAVAGH**S**MGGGGTLRIAEQNGLPVALAFTPWEPGPPRGGLLLLLLGLGGG**D**TVAPVSQHAIPFYQNSGLGLALP**H**FAPNNGRAFEALEELLAQCQ

**> P4**

SNALLLLLLLLLLLGGGLLLALAGGYTADASSLAWLGRRLASLFGVLLLSGGYPDSRASQLSAALNYLRTQLASLGLDANRLAVAGH**S**MGGGGTLRIAEQNRGGLLVLVFTPWESGPARGGRLVVLVVGAS**D**TVAPVSQHAIPFYQNLRGGLVVLVGLP**H**FAPNRGSPELLEALLRLLRLLLQCQ

**> P5**

SNLLLLLLLGAGALALYAGYTADASSLAWLGRRLASLGLAVLLLSGGGYPDSRASQLSAA
LNYLRTQLASLGLDANRLAVAGH**S**MGGGGTLRIAEQNRGGLLALAFTPWEGGRAGRAVL
VLGS**D**TVAPVSQHAIPFYQNARPGGGLLLLLGGN**H**FAPNNGRALDALEELLQLLQQCQ

**> P6**

SNPRPRLLALASGYTADASSLAWLGRRLASLFPNLGVALVDGRNEYPDSRASQLSAALNY
LRTLAEAQGLGLDANRLAVAGH**S**MGGGGTLRIAEQNGVPFVVFTPWDAEPPRGGRLLLV
VGGRN**D**TVAPVSQHAIPFYQNNPNLRLVLLPGN**H**FAPNDPELLEELLELLLELLLLLPQCQ

**> P7**

SNCSLLLLLLSDLEELLELLEALLGGLAFALASGYTADASSLAWLGRRLASLLVNGNGYPD
SRASQLSAALNYLRTNRPGLDANRLAVAGH**S**MGGGGTLRIAEQNGLPLALLLTPWGGGG
GGGGLLGGG**D**TVAPVSQHAIPFYQNLPGGRLLLVLGG**H**FAPNNGRGGLVLLLLRCQ

**> P8**

SNCLLLLSLEEALARLAARGLAVALASGYTADASSLAWLGRRLASLGVGVLLLSAGYPDS
RASQLSAALNYLRTNLASRLGLDANRLAVAGH**S**MGGGGTLRIAEQNGVDALLFTPWGGG
GGGG**D**TVAPVSQHAIPFYQNRGLLVVAGN**H**FAPNNGLLLGGLLLLLLRCQ

**> P9**

SNLAALLSRLGIAIGSGYTADASSLAWLGRRLASLLGVLLVNGNGYPDSRASQLSAALNY
LRTNRPSLDANRLAVAGH**S**MGGGGTLRIAEQNRGGLLAVVFTPWGGGGGGGLLLLLAG
**D**TVAPVSQHAIPFYQNGLLLLP**H**FAPNDPADADELLALLLRLLLELLALLEACQ

**> P10**

SNLALLLLLLRGLALALAAGYTADASSLAWLGRRLASGGVLVVGGGYPDSRASQLSAAL
NYLRTNGLRLAGLPLDANRLAVAGH**S**MGGGGTLRIAEQNPGLDLALVLTPWGGGGSG**D**T
VAPVSQHAIPFYQNRGLLLAAP**H**FAPNNGLALELLLELLAALLGGGGLLLLLLLDGGGGV
VLLLLNCQ

**Supplementary Table 3: Changes in the hydrophilic and hydrophobic properties and gyration radii of the proteins after sequence refinement using iterative $RF_{joint}$ and ProteinMPNN.**

| Protein | Hydrophilic surface area ($nm^2$) | Hydrophobic surface area ($nm^2$) | Total surface area ($nm^2$) | Hydrophobic surface / Total surface | $R_g$ (nm) |
|---|---|---|---|---|---|
| P4 | 45.0 | 50.5 | 95.5 | 52.9% | 1.58 |
| P4-a | 46.9 | 40.7 | 87.6 | 46.5% | 1.59 |
| P4-a-1 | 36.9 | 42.0 | 78.9 | 53.3% | 1.53 |
| P4-a-2 | 36.4 | 43.6 | 80.0 | 54.5% | 1.55 |
| P5 | 45.5 | 41.9 | 87.4 | 47.9% | 1.51 |
| P5-a | 46.9 | 35.5 | 82.3 | 43.1% | 1.50 |
| P5-a-1 | 40.8 | 39.7 | 80.5 | 49.3% | 1.49 |
| P5-a-2 | 43.4 | 37.9 | 81.3 | 46.6% | 1.50 |
| P7 | 45.1 | 39.3 | 84.3 | 46.6% | 1.49 |
| P7-a | 47.4 | 37.4 | 84.8 | 44.1% | 1.53 |

**References:**


1. Madeira, F., et al. Search and sequence analysis tools services from EMBL-EBI in 2022. *Nucleic Acids Res.* **50**, W276-W279 (2022).

2. Flachsenberg, F., et al. A Consistent Scheme for Gradient-Based Optimization of Protein-Ligand Poses. *J. Chem. Inf. Model* **60**, 6502–6522 (2020).
